# Behavioral changes and growth deficits in a CRISPR engineered mouse model of the schizophrenia-associated 3q29 deletion

**DOI:** 10.1101/479949

**Authors:** Timothy P. Rutkowski, Ryan H. Purcell, Rebecca M. Pollak, Stephanie M. Grewenow, Georgette M. Gafford, Tamika Malone, Uswa A. Khan, Jason P. Schroeder, Michael P. Epstein, Gary J. Bassell, Stephen T. Warren, David Weinshenker, Tamara Caspary, Jennifer Gladys Mulle

**Affiliations:** Department of Human Genetics, Emory University School of Medicine, Atlanta, GA 30322; Department of Cell Biology, Emory University School of Medicine, Atlanta, GA 30322

## Abstract

The 3q29 deletion confers increased risk for neuropsychiatric phenotypes including intellectual disability, autism spectrum disorder, generalized anxiety disorder, and a >40-fold increased risk for schizophrenia. To investigate consequences of the 3q29 deletion in an experimental system, we used CRISPR/Cas9 technology to introduce a heterozygous deletion into the syntenic interval on C57BL/6 mouse chromosome 16. mRNA abundance for 20 of the 21 genes in the interval was reduced by ~50%, while protein levels were reduced for only a subset of these, suggesting a compensatory mechanism. Mice harboring the deletion manifested behavioral impairments in multiple domains including social interaction, cognitive function, acoustic startle, and amphetamine sensitivity, with some sex-dependent manifestations. Additionally, 3q29 deletion mice showed reduced body weight throughout development consistent with the phenotype of 3q29 deletion syndrome patients. Of the genes within the interval, *DLG1* has been hypothesized as a contributor to the neuropsychiatric phenotypes. However, we show that *Dlg1*^+/-^ mice did not exhibit the behavioral deficits seen in mice harboring the full 3q29 deletion. These data demonstrate the following: the 3q29 deletion mice are a valuable experimental system that can be used to interrogate the biology of 3q29 deletion syndrome; behavioral manifestations of the 3q29 deletion may have sex-dependent effects; and mouse-specific behavior phenotypes associated with the 3q29 deletion are not solely due to haploinsufficiency of *Dlg1*.

## Introduction

3q29 deletion syndrome is caused by heterozygous deletion of a 1.6 Mb interval containing 21 protein coding genes. Individuals with the deletion are at increased risk for intellectual disability, autism spectrum disorder (ASD), generalized anxiety disorder, and schizophrenia (SZ) (1–7). In the largest association study of copy number variation and SZ to date, the 3q29 deletion emerges as one of only 8 loci achieving genome-wide significance (3). Separate analyses have shown that the 3q29 deletion confers an exceptional risk for SZ, with an estimated effect size of >40 (8, 9). This large effect size, coupled with the relatively low complexity of the interval and the high synteny between the human and mouse genomes, make this variant an ideal candidate for the development of a mouse experimental system. This is further aided by CRISPR/Cas9 technology facilitating engineered genomic rearrangements (10). Collectively, the 3q29 deletion is a compelling region to study because it provides an opportunity for genetic dissection of complex neuropsychiatric phenotypes.

Of the 21 genes within the interval, *DLG1* and *PAK2* have received attention as attractive candidates contributing to neuropsychiatric phenotypes (11). DLG1 (a.k.a. SAP97) is a scaffolding protein that interacts with N-methyl-D-aspartate (NMDA) and AMPA type glutamate receptors (12, 13). Suggestive evidence for association between *DLG1* and neuropsychiatric phenotypes comes from sequencing studies where SZ cases are enriched for *DLG1* variants compared to controls (14, 15). An expression study of postmortem tissue found decreased DLG1 in the prefrontal cortex in SZ patients (16). PAK2 is implicated by its function as a regulator of cytoskeletal dynamics. Based on the current evidence for their possible involvement in neuropsychiatric phenotypes, mouse models have been created for both *Dlg1* and *Pak2* deficiency. A brain-specific conditional *Dlg1-* deletion mouse displays some sex-specific behavioral phenotypes, including subtle cognitive and motor deficits (17). A *Pak2*^+/-^ mouse model displays autism-related behaviors and neuronal deficits (18). Drosophila models also support involvement of *dlg* and *pak*, but behavioral and molecular phenotypes are only seen when *dlg* and *pak* are simultaneously reduced (19).

We undertook a two-step approach to better understand the neuropsychiatric consequences of the 3q29 deletion and the possible contribution of *DLG1* to these phenotypes. First, we created a mouse harboring a heterozygous 3q29 deletion (henceforth referred to as B6.Del16^+/*Bdh1-Tfrc*^ mice [nomenclature adapted from Mouse Genome Informatics, MGI:6241487]), and assessed gene expression, protein abundance, developmental weight trajectories, and behavioral changes associated with the deletion. Second, we used *Dlg1*^+/-^ mice to test the hypothesis that *DLG1* haploinsufficiency alone is sufficient to manifest B6.Del16^+/*Bdh1-Tfrc*^-associated phenotypes. We performed parallel analyses in both B6.Del16^+/*Bdh1-Tfrc*^ and *Dlg1*^+/-^ mice. Our results allow for integration of existing data from mouse models of other 3q29 interval genes and pave the way for careful and rigorous dissection of the genetic drivers for neuropsychiatric phenotypes within the 3q29 interval.

## Materials and Methods *(see Supplemental Material for detailed Methods and Protocols)*

### Mouse strains and alleles

All mouse work was performed under the approved guidelines of the Emory University IACUC. To generate the 3q29 deletion in the mouse, two CRISPR gRNAs were designed at the syntenic loci in the mouse genome. The Emory Transgenic and Gene Targeting core injected 50ng/ml of each gRNAs and 100ng/ml Cas9 RNA into single-cell C57BL/6N zygotes. Embryos were cultured overnight and transferred to pseudopregnant females. Resulting pups were screened for the deletion via PCR and for an absence of genomic rearrangemnets by Southern blot (See **Supplemental Materials** for primer sequences used for genotyping and probe details). The founders used (#127, #131) were backcrossed to C57BL/6N and analysis commenced in the N4 generation. All mice were maintained on a C57BL/6N background sourced from Charles River Laboratories. The *Dlg1*^+/-^ mice [MGI:3699270] (20) were obtained from Dr. Jeffrey Miner (Washington University in St Louis) on a mixed 129/C57BL/6J background. The mice were backcrossed using marker-assisted breeding (DartMouse^™^, https://dartmouse.org) to obtain N6 Dlg1^+/-^ mice that were 99% congenic on a C57BL/6N background (henceforth referred to as B6.*Dlg1*^+/-^). Number of animals used in experiments is indicated in figure legends.

### Gene and protein expression in mouse brain tissue

RNA was isolated from forebrain samples from 16-20 week-old mice and gene expression was measured by real-time quantitative PCR. See **Supplemental Table 1** for list of Taqman assay IDs. Protein was isolated from forebrain samples from 16-20 week-old mice according to standard procedures and protein levels were measured by Western blot. See **Supplemental Table 2** for antibodies.

### Behavior Tests

Mice were on a 12-hour light/dark cycle and were given food and water *ad libitum*. Mice were between the ages of 16-20 weeks when behavioral testing commenced. All equipment was cleaned using Virkon. No method of randomization was utilized in the behavior studies.

#### Social Interaction and Morris water maze

The three-chambered social interaction paradigm was adapted from Yang et al. (21). The Morris water maze was conducted using the same paradigm as Chalermpalanupap et al. (22).

#### Prepulse Inhibition (PPI)

The San Diego Instruments (La Jolla, CA) SR-LAB startle response system was used to perform PPI. We used a two-day paradigm. On day one, we tested the ability of the mouse to startle to a series of increasing tones. On day two, we subjected each mouse through the prepulse inhibition paradigm. PPI was calculated as a percentage using the following equation: ((startle-startle.PP/startle))*100.

#### Amphetamine induced locomotor activity

The assay was performed in a locomotor chamber (San Diego Instruments), which consisted of a plexiglass cage (48×25×22cm) containing corncob bedding that rested between an apparatus containing infrared beams. The mice were given an injection of either saline or amphetamine (2.5 or 7.5 mg/kg, i.p.) amphetamine, and post-injection ambulations were measured for 2 hours. All treatments were spread over 3 weeks and were counter-balanced such that not all of the mice received the same injection in a given week. For the 7.5mg/kg amphetamine dose, a subset of mice was video recorded between the 30-60 minute post-injection time point. These videos were scored for stereotypy using the criteria as described previously (23) by an experimenter blinded to genotype.

#### Additional Measures

Details on paradigms for circadian rhythm, elevated plus maze, open field, marble burying, social interaction, prepulse inhibition, fear conditioning, and histology can be found in *Supplemental Materials*.

Sample sizes of 8-16 mice per sex and genotype were used in agreement with existing literature.

### Analysis

#### Gene expression

Data were analyzed by two-way, repeated measures ANOVA followed by multiple comparisons with Sidak’s correction, where appropriate (i.e. only when a significant [p<0.05] genotype effect or interaction was detected).

#### Behavioral Tests

All behavior tests were analyzed following these parameters unless otherwise specified. Behavior was analyzed by unpaired t-test (when comparing 2 groups) or two-way, repeated measures ANOVA (when comparing more than 2 groups). followed by multiple comparisons with Sidak’s correction where appropriate (i.e. only when a significant [p<0.05] genotype effect or interaction was detected).

#### Amphetamine-induced Locomoter Activity

All analyses performed in R (24). Because the ambulation data were not normally distributed, the inverse normal function was used to transform the data to an approximately normal distribution. Proper transformation of the data was confirmed with the Shapiro-Wilk normality test implemented using the stats package (24). Linear mixed-effects models with restricted maximum likelihood estimation were implemented using the lme4 package (25). All models included subject ID and timepoint as random effects; males and females were analyzed separately. Models tested for the main effects of genotype and treatment (saline, 2.5 mg/kg amphetamine, and 7.5 mg/kg amphetamine), and for an interaction between genotype and treatment. Saline was set as the reference treatment and WT was set as the reference genotype for all analyses. P values were calculated using Satterthwaite’s method using the lmerTest package (26).

#### Acoustic startle and Growth Curves

Analyses for acoustic startle and growth curves were performed in R (24). Additional details on the analyses for these tasks can be found in the supplemental materials.

All data represented as mean ± SEM.

## Results

### Confirmation of deletion using CRISPR/Cas9

To generate mice that recapitulated the 3q29 deletion, we took advantage of several features of the syntenic region on mouse chromosome 16. The mouse and human regions are almost identical with all 21 genes present in the same order (*Bex6* present in the mouse, not human). The mouse syntenic region is inverted; inversion breakpoints are identical to the SZ and ID-associated deletion breakpoints. The mouse region is also slightly smaller compared to the human region, 1.26 MB vs. 1.6 MB (**Figure 1a**). To generate B6.Del16^+/*Bdh1-Tfrc*^ mice, we mimicked the 3q29 deletion breakpoints by designing CRISPR gRNAs proximal to *Bdh1* and distal to *Tfrc* (**Figure 1b**). We performed PCR to confirm the presence of the heterozygous deletion in potential founder animals. To investigate whether the double strand breaks during CRISPR/Cas9 editing created additional genomic rearrangements in the region, we performed a Southern blot using a probe centromeric to the proximal breakpoint (**Figure 1b**). The radiolabelled probe hybridized to a wild-type 5.2kb XbaI fragment on the intact, non-deleted chromosome and a 6.2 kb XbaI fragment on the deleted chromosome. Of the 23 potential founders, we identified 7 (30%) animals that appeared to be properly targeted. The PCR products across the deletion were sequenced to determine the precise breakpoints for the two founders used [#127 and #131] (See **Supplemental Materials**).

**Figure 1:**
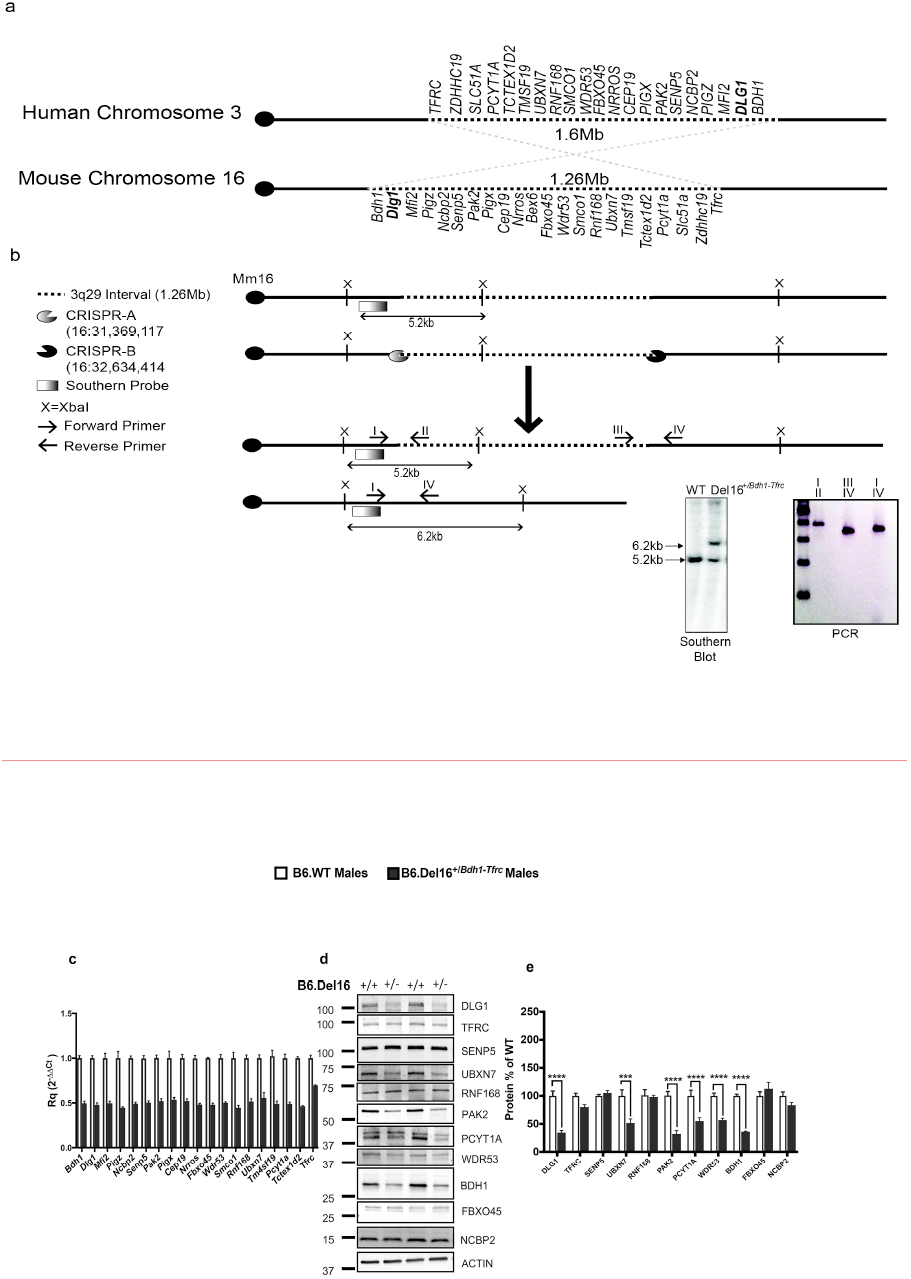
Generation of 3q29 deletion on the C57BL/6N background using CRISPR/Cas9. **(a)** The 3q29 region is located on human chromosome 3 [recurrent deletion coordinates (GRCh38.p12)-Chr3:195998129-197623129], and the syntenic 3q29 region is located on mouse chromosome 16 [mouse 3q29 deletion coordinates (GRCm38.p3)-Chr16:31369117-32634414]. **(b)** Two CRISPRs were designed (CRISPR-A [Chr.16:31369117] and CRISPR-B [Chr.16:32634414]) to create a heterozygous 3q29 deletion. The deletion was confirmed using southern blots (5.2kb=wild-type band, 6.2kb=deletion band) and PCR. **(c)** B6.Del16^+/*Bdh1-Tfrc*^ male mice have ~50% decreased gene expression in most of the genes within the 3q29 interval (with the exception of *Tfrc*). All genes are significantly reduced (p<0.0001). **(d)** Protein expression analyses reveals a significant reduction in DLG1, UBXN7, PAK2, PCYT1A, WDR53, and BDH1 (***p<0.0005, ****p<0.0001).

We analyzed the expression of the 19/22 genes (*Slc51a* and *Zdhhc19* are not expressed in brain; *Bex6* is mouse-specific and not present in the human interval) using forebrain tissue from both adult female and male B6.Del16^+/*Bdh1-Tfrc*^ mice. Using real-time PCR (RT-PCR), we found that expression of 18/19 genes tested was 50% reduced in female (**Supplemental Figure 1a**) and male (**Figure 1c**) B6.Del16^+/*Bdh1-Tfrc*^ mice compared to B6.WT littermates. The exception was *Tfrc*, which was not significantly reduced in female B6.Del16^+/*Bdh1-Tfrc*^ mice, and only 30% reduced in male B6.Del16^+/*Bdh1-Tfrc*^ mice. We then examined protein expression of eleven of the highest-expressed genes by Western blot. In both females (**Supplemental Figure 1b-c**) and males (**Figure 1d-e**), we found that only 6 of 11 proteins (DLG1, UBXN7, PAK2, PCYT1A, WDR53, and BDH1) were found to be significantly reduced in B6.Del16^+/*Bdh1-Tfrc*^ mouse forebrain tissue compared to B6.WT littermates; no genotype differences were detected for TFRC, SENP5, RNF168, FBXO45, and NCBP2. Thus, while the transcripts for 18 genes in the interval have decreased expression in a manner consistent with their haploinsufficiency, the corresponding protein levels are not always changed in B6.Del16^+/*Bdh1-Tfrc*^ mice. These data demonstrate that B6.Del16^+/*Bdh1-Tfrc*^ mice recapitulate the genetic lesion <u>in</u> 3q29 deletion syndrome and suggest leads for phenotypic driver genes.

### B6.Del16^+/*Bdh1-Tfrc*^ display growth deficits

Individuals with the 3q29 deletion demonstrate reduced weight at birth, and this may persist throughout childhood (2). To assess whether the B6.Del16^+/*Bdh1-Tfrc*^ mice display a growth phenotype, we generated cohorts of female and male B6.Del16^+/*Bdh1-Tfrc*^ mice along with wild-type (B6.WT) littermate controls. We weighed mice weekly over 16 weeks (starting at P8) and found that female B6.Del16^+/*Bdh1-Tfrc*^ mice weighed on average 2.24 g less than WT littermates (p<0.0001) and male B6.Del16^+/*Bdh1-Tfrc*^ weighed on average 1.61 g less than WT littermates (p<0.0005, **Figure 2a-b**). These data indicate that the B6.Del16^+/*Bdh1-Tfrc*^ mice recapitulate the reduced growth observed in study subjects with the 3q29 deletion.

**Figure 2:**
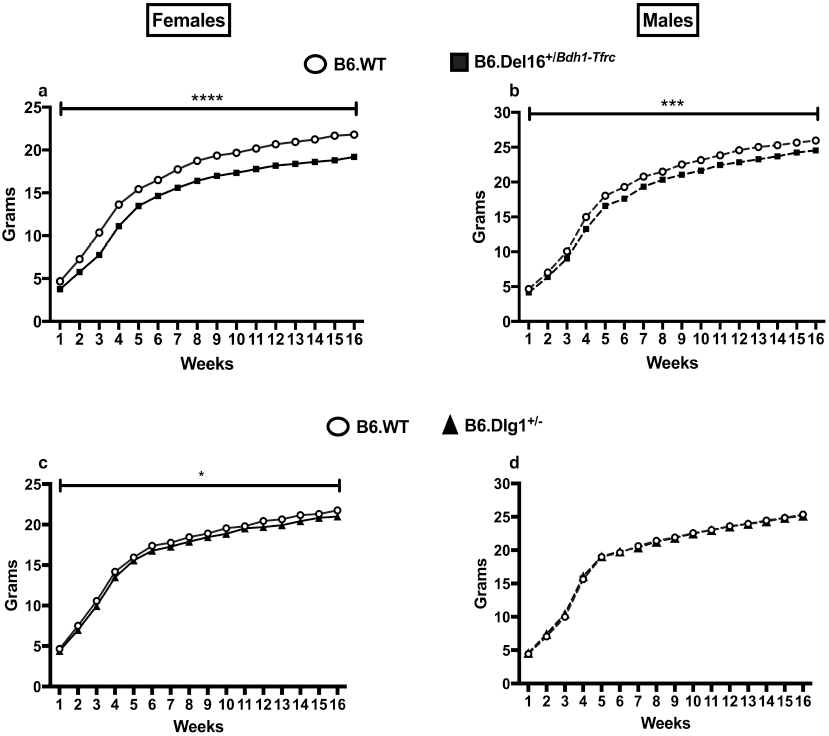
Growth deficits in B6.Del16^+/*Bdh1-Tfrc*^ mice and female B6.*Dlg1*^+/-^ mice. **(a)** Female (****p<0.0001) [N=34 wild-type, 32 mutant] and **(b)** male (***p<0.0005) [N=33 wild-type, 27 mutant] B6.Del16^+/*Bdh1-Tfrc*^ mice weigh significantly less by genotype compared to B6.WT littermates. **(c)** Female (*p<0.05) [N=23 wild-type, 25 mutant], but not **(d)** male (p>0.05) [N=36 wild-type, 30 mutant], B6.*Dlg1*^+/-^ mice weigh significantly less compared to their B6.WT littermates. Weight data analyzed as outlined in methods.

We then assessed B6.*Dlg1*^+/-^ mice for weight deficits over 16 weeks (starting at P8). Female B6.*Dlg1*^+/-^ mice weigh an estimated 0.78 g less than WT (p<0.05). The weight effect was not seen in male B6.*Dlg1*^+/-^ mice (p>0.05, **Figure 2c-d**). The effect sizes for the orthologous 3q29 deletion are significantly larger than the effect sizes for the *Dlg1* heterozygote for females (p<0.0005). Taken together, these data indicate that the growth phenotype in B6.Del16^+/*Bdh1-Tfrc*^ mice is not due to heterozygosity of *Dlg1* alone.

### B6.Del16^+/*Bdh1-Tfrc*^ mice have smaller brains, but normal brain architecture

We examined whole brains in B6.Del16^+/*Bdh1-Tfrc*^ mice for gross structural or anatomical changes. Both female (t_21_=7.316, p<0.0001) and male (t_15_=5.231, p<0.0005) B6.Del16^+/*Bdh1-Tfrc*^ brains are *smaller* compared to B6.WT littermates (**Supplemental Figure 2a**). However, when we normalized the brain weight to body weight, we found that the brain:body weight ratio was *increased* in female (t_21_=3.876, p<0.001), but not male (t_15_=0.4154, p>0.05), B6.Del16^+/*Bdh1-Tfrc*^ brains compared to B6.WT littermates (**Supplemental Figure 2b**). We analyzed adult coronal sections with a cresyl violet stain and saw no gross differences in B6.Del16^+/*Bdh1-Tfrc*^ mice compared to their B6.WT littermates for either sex (**Supplemental Figure 2c-d**). We assessed for any perturbation in the cortical plate using an antibody for T-box, Brain1 (TBR1). In E15.5 embryos, B6.Del16^+/*Bdh1-Tfrc*^ mice appeared to specify the cortical plate normally (**Supplemental Figure 3**). Collectively, these data revealed altered brain size but no obvious architectural phenotypes in B6.Del16^+/*Bdh1-Tfrc*^ brain morphology.

### B6.Del16^+/*Bdh1-Tfrc*^ and B6.Dlg1^+/-^ have normal locomotion

We assessed locomotion in B6.Del16^+/Bdh1-Tfrc^ and B6.Dlg1^+/-^ mice using automated activity chambers by recording the number of ambulations within the chambers. We found no differences in ambulations immediately following exposure to a novel environment or over the ensuing 24h in either female (main effect of genotype, *F_1,30_*=2.387, p>0.05) or male (main effect of genotype, *F_1,28_*=0.3563, p>0.05) B6.Del16^+/*Bdh1-Tfrc*^ mice (**Supplemental Figure 4a-b**). Similarly, B6.*Dlg1*^+/-^ females (main effect of genotype, *F_1,15_*=0.3667, p>0.05) or males (main effect of genotype, *F_1,11_*=0.3087, p>0.05) (**Supplemental Figure 4c-d**) also showed no differences in ambulations compared to their respective B6.WT littermates. Thus, neither B6.Del16^+/*Bdh1-Tfrc*^ nor B6.*Dlg1*^+/-^ mice have locomotor deficits, suggesting that behavioral phenotypes can be assessed, as the mice do not exhibit an obvious movement confound.

### B6.Del16^+/*Bdh1-Tfrc*^ male mice have social interaction deficits

Because ASD is one of the phenotypes reported in 3q29 deletion patients (2, 27), we tested the mice for social interaction deficits using the three-chamber social approach task. We measured duration of olfactory investigation to gauge how much a given subject mouse was interacting either with the empty cup or with the cup containing the stranger mouse. As expected, both female (t_26_=7.176, p<0.0001) and male (t_28_=4.018, p<0.0005) B6.WT interacted significantly more with the stranger mouse compared to the empty cup. Female (t_24_=3.237, p<0.005) B6.Del16^+/*Bdh1-Tfrc*^ also showed normal sociality. By contrast, male (t_28_=1.769, p=0.0878) B6.Del16^+/*Bdh1-Tfr*^ mice exhibited abnormal sociality as they did not show a significant preference for the stranger mouse over the empty cup (**Figure 3a**). We then tested female and male B6.*Dlg1*^+/-^ mice to determine whether haploinsufficiency of *Dlg1* led to social impairment (**Figure 3b**). We did not observe any deficits in either female (t_24_=4.716, p<0.0001) or male (t_24_=2.846, p<0.01) B6.*Dlg1*^+/-^ mice or their B6.WT littermates [female (t_26_=3.711, p<0.005), (male t_22_=6.29, p<0.0001)]. These data indicate that male B6.Del16^+/*Bdh1-Tfr*^ mice display social impairment that is not solely attributable to *Dlg1* haploinsufficiency.

**Figure 3:**
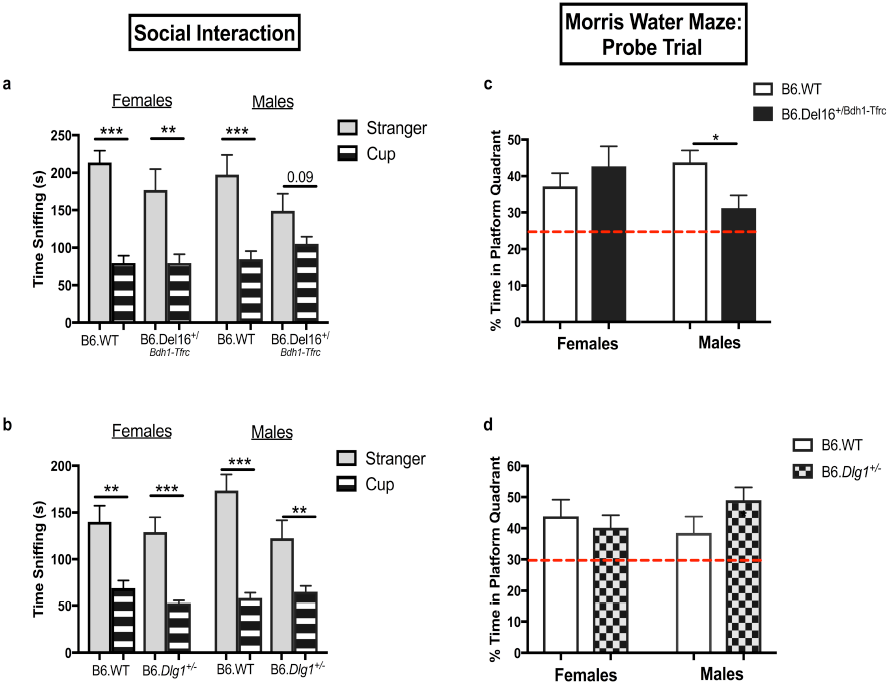
Social and cognitive impairment in male B6.Del16^+/*Bdh1-Tfrc*^ mice. **(a)** Female B6.Del16^*+/Bdh1-Tfrc*^ mice interact significantly more with the stranger under the cup compared to empty cup alone (**p<0.01). similar to their B6.WT littermates (***p<0.0005) [N=14 wild-type, 14 mutant]. Male B6.Del16^*+/Bdh1-Tfrc*^ mice do not show a significant preference for interacting with either the stranger under the cup or the empty cup (p=0.09). Male B6.WT mice significantly prefer to interact with the stranger under the cup compared to the empty cup (***p<0.0005) [N=15 wild-type, 15 mutant]. **(b)** Female B6.*Dlg1*^+/-^ mice interact significantly more with the stranger under the cup compared to empty cup alone (***p<0.0005)) similar to their B6.WT littermates (**p<0.01)) [N=14 wild-type, 13 mutant]. Male B6.*Dlg1*^+/-^ mice interact significantly more with the stranger under the cup compared to the empty cup (**p<0.01) similar to their B6.WT littermates (***p<0.0005) [N=12 wild-type, 13 mutant]. **(c)** Female B6.Del16^+/*Bdh1-Tfrc*^ mice spend a similar amount of time in the platform quadrant compared to B6.WT littermates (p>0.05) [N=16 wild-type, 16 mutant]. Male B6.Del16^*+/Bdh1-Tfrc*^ mice spend significantly less time in the platform quadrant compared to B6.WT littermates (*p<0.05) [N=15 wild-type, 15 mutant]. **(d)** Female (p>0.05) [8 wild-type, 11 mutant] and male (p>0.05) [9 wild-type, 9 mutant] B6.*Dlg1*^+/-^ mice spend a similar time in the platform quadrant compared to their B6.WT littermates. The dashed red line denotes 25%/chance. All data analyzed by two-tailed Students t-test. Results represent the mean ± SEM.

### B6.Del16^+/*Bdh1-Tfrc*^ male mice have spatial memory deficits

In the Morris water maze (MWM), female B6.Del16^+/*Bdh1-Tfrc*^ mice showed a similar pattern of learning compared to their B6.WT littermates as there was no difference in latency (main effect of genotype, *F_1,30_*=0.07352, p>0.05), or distance (main effect of genotype, *F_1,30_*=0.6494, p>0.05) to find the hidden platform during the training portion of the MWM. While there was no genotype difference in swim speed (main effect of genotype, *F_1,30_*=2.188, p>0.05), there was a significant interaction (genotype x time, *F_4,120_*, p<0.005). Sidak’s post-hoc analysis revealed female B6.Del16^+/*Bdh1-Tfrc*^ mice swim faster on day 5 compared to B6.WT littermates (p<0.005) (**Supplemental Figure 5a-c**). Male B6.Del16^+/*Bdh1-Tfrc*^ mice showed a trend towards increased latency that did not reach significance (main effect of genotype, *F_1,28_*=3.922, p=0.057), swam a greater distance (main effect of genotype, *F_1,28_*=4.621, p<0.05), but had similar swim speed (main effect of genotype, *F_1,28_*=0.493, p>0.05) to reach the hidden platform on day 5 compared to B6.WT littermates (**Supplemental Figure 5d-f**). We then tested for spatial memory deficits in the probe trial (**Figure 3c**) and found no difference in percentage of time spent in the quadrant that formerly contained the platform between female B6.Del16^+/*Bdh1-Tfrc*^ mice and their B6.WT littermates (t_30_=0.8357, p>0.05). However, male B6.Del16^+/*Bdh1-Tfrc*^ mice spent significantly less time in the platform quadrant compared to their B6.WT littermates (t_28_=2.592, p<0.05).

We tested both female and male B6.*Dlg1*^+/-^ mice in the MWM. There were no significant differences in latency (main effect of genotype, *F_1,17_*=0.1776, p>0.05), distance (main effect of genotype, *F_1,17_*=0.1407, p>0.05), or swim speed (main effect of genotype, *F_1,17_*=0.5809, p>0.05) in female **(Supplemental Figure 6a-c)** or male [latency: (main effect of genotype, *F_1,16_*=0.2816, p>0.05), distance: (main effect of genotype, F_1,16_=0.3065, p>0.05), speed: (main effect of genotype, *F_1,16_*=0.0031, p>0.05) **(Supplemental Figure 6d-f)** B6.*Dlg1*^+/-^ mice compared to B6.WT littermates. During the probe trial **(Figure 3d)**, both female (t_17_=0.5685, p>0.05) and male (t_16_=1.573, p>0.05) B6.*Dlg1*^+/-^ mice spent a similar amount of time in the platform quadrant compared to B6.WT littermates. Collectively, these data show male B6.Del16^+/*Bdh1-Tfrc*^ mice exhibit spatial learning and memory deficits that are not solely due to haploinsufficiency of *Dlg1*.

### B6.Del16^+/*Bdh1-Tfrc*^ mice display increased startle response

We examined potential differences in response to an acoustic startle and in sensorimotor gating using PPI. Female B6.Del16^+/*Bdh1-Tfrc*^ mice **(Figure 4a)** displayed a significant increase in acoustic startle response compared to their B6.WT littermates (p < 0.05). Male B6.Del16^+/*Bdh1-Tfrc*^ mice showed a similar trend of increased startle, but it did not reach significance (p = 0.056) **(Figure 4b)**. Because the magnitude of the measured startle response can be affected by weight, we performed statistical analyses to correct for the decreased weight of B6.Del16^+/*Bdh1-Tfrc*^ mice. Our analyses revealed that weight did not contribute significantly to startle response (p > 0.25), indicating that the weight deficits observed in B6.Del16^+/*Bdh1-Tfrc*^ mice did not affect the measured startle response. Female **(Figure 4c)** (main effect of genotype, *F_1,17_*=0.6714, p>0.05) and male **(Figure 4d)** (main effect of genotype, *F_1,16_*=0.4238, p>0.05) B6.*Dlg1*^+/-^ mice displayed normal acoustic startle. We assessed for deficits in sensorimotor gating using PPI. Female B6.Del16^+/*Bdh1-Tfrc*^ mice had similar PPI compared to B6.WT littermates (main effect of genotype, F_*1,30*_=0.083), but did have a significant interaction (genotype x prepulse, *F_3,90_*=3.527, p<0.05). Sidak’s post-hoc analysis revealed no significant differences at the respective prepulses (p>0.05). Male B6.Del16^+/*Bdh1-Tfrc*^ mice did not show any differences compared to B6.WT littermates (main effect of genotype, *F_1,28_*=0.1715, p>0.05). Furthermore, neither female (main effect of genotype, *F_1,17_*=1.654, p>0.05) nor male (main effect of genotype, *F_1,16_*=1.998, p>0.05) B6.*Dlg1*^+/-^ mice showed any differences in PPI compared to their B6.WT littermates **(Figure 4e-h)**. These data indicate that mice harboring the 3q29 deletion have an increased response to an acoustic startle but mostly normal sensorimotor gating, whereas happloinsufficiency of *Dlg1* does not alter either measure.

**Figure 4:**
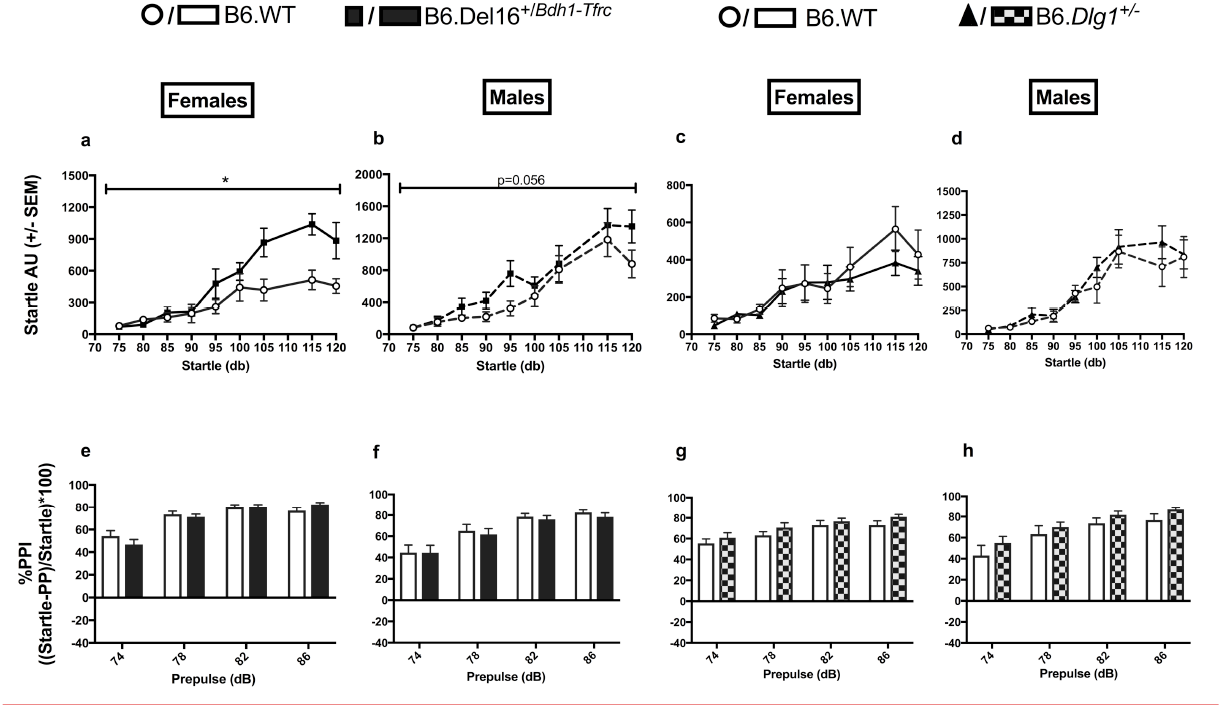
Increased startle, but normal prepulse inhibition, in B6.Del16^+/*Bdh1-Tfrc*^ mice. **(a)** Female [N=16 wild-type, 16 mutant] and **(b)** male [N=15 wild-type, 15 mutant] B6.Del16^*+/Bdh1-Tfrc*^ mice startle more compared to B6.WT littermates. **(c)** Female [N=8 wild-type, 11 mutant] and **(d)** male [N=9 wild-type, 9 mutant] B6.*Dlg1*^+/-^ mice startle similarly to B6.WT littermates. **(e)** Female [N=16 wild-type, 16 mutant] **(f)** male [N=15 wild-type, 15 mutant] B6.Del16^+/*Bdh1-Tfrc*^ mice have similar prepulse inhibition compared to B6.WT littermates. **(g)** Female (*F_1,17_*=1.654) and (*h*) male (*F_1,16_*=1.998) B6.*Dlg1*^+/-^ mice have similar prepulse inhibition compared to B6.WT littermates. Startle data analyzed as outlined in methods. PPI data analyzed by two-way, repeated measures Anova followed by multiple comparisons. Results represent the mean ± SEM (*p<0.05)

### Amphetamine-induced locomotor activity is attenuated in B6.Del16^+/*Bdh1-Tfrc*^ mice

Chronic amphetamine use can lead to SZ symptoms, and amphetamine can exacerbate SZ symptoms in individuals that already have the disorder (28). Because of the strong association between the 3q29 deletion and SZ, we tested amphetamine sensitivity in both B6.Del16^+/*Bdh1-Tfrc*^ and B6.*Dlg1*^+/-^mice. Following saline administration, neither female nor male B6.Del16^+/*Bdh1-Tfrc*^ mice displayed significant differences in ambulatory activity compared to their B6.WT littermates (main effect of genotype, p>0.05). At both the 2.5mg/kg and 7.5mg/kg doses of amphetamine, both female and male B6.WT mice were significantly more active compared to the respective B6.WT mice that received saline (main effect of treatment, p<0.0001). Amphetamine-induced locomotion was attenuated in female B6.Del16^+/*Bdh1-Tfrc*^ mice following the 2.5mg/kg (treatment x genotype interaction, p<0.005) and 7.5mg/kg (treatment x genotype interaction, p<0.0001) doses compared to B6.WT littermates **(Figure 5a)**. Male B6.Del16^+/*Bdh1-Tfrc*^ mice did not show any differences in activity following the 2.5mg/kg dose (treatment x genotype interaction, p>0.05), but did show a reduction in activity following the 7.5mg/kg dose (treatment x genotype interaction, p<0.0001) of amphetamine compared to B6.WT littermates **(Figure 5b)**. Because stereotypy can occlude horizontal locomotion at high doses of amphetamine, we video recorded mice from the 35-65 min post-injection and manually scored the videos for stereotypy as previously reported (23), but did not observe any significant differences (data not shown). We then assessed amphetamine sensitivity in B6.*Dlg1*^+/-^ mice. Neither female nor male B6.*Dlg1*^+/-^ mice displayed significant differences in ambulatory activity compared to their B6.WT littermates following administration of saline (main effect of genotype, p>0.05). At both the 2.5mg/kg and 7.5mg/kg dose of amphetamine, both female and male B6.WT mice were significantly more active compared to the respective B6.WT mice that received saline (main effect of treatment, p<0.0001). Female B6.*Dlg1*^+/-^ mice did not display differences in activity at the 2.5mg/kg (treatment x genotype interaction, p>0.05) or 7.5mg/kg (treatment x genotype interaction, p>0.05) amphetamine doses compared to their B6.WT littermates **(Figure 5c)**. Male B6.*Dlg1*^+/-^ mice showed a small but significant increase in ambulatory activity following administration of 2.5mg/kg amphetamine (treatment x genotype interaction, p<0.001), but no significant changes in activity following the 7.5mg/kg dose (treatment x genotype interaction, p>0.05) compared to B6.WT littermates **(Figure 5d)**. Taken together, these data show that B6.Del16^+/*Bdh1-Tfrc*^ mice display attenuated amphetamine-induced locomotor activity that is not solely due to haploinsufficiency of *Dlg1*.

**Figure 5:**
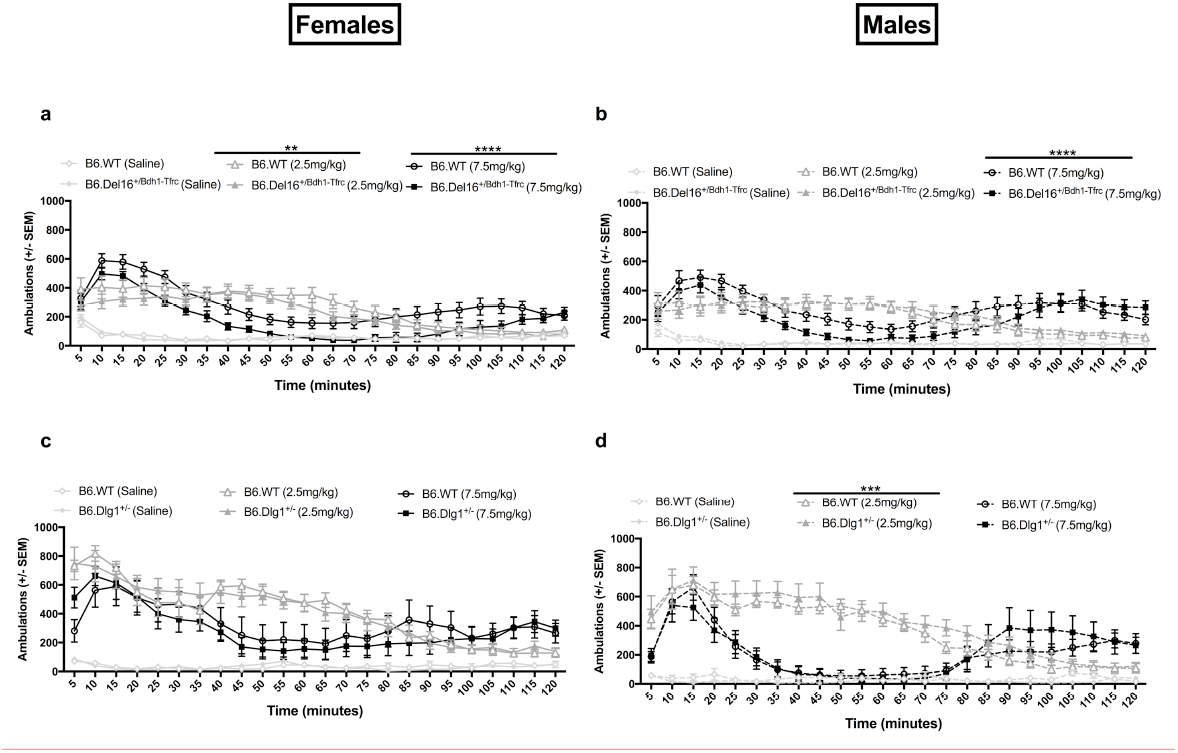
Attenuated activity after a high dose of amphetamine in B6.Del16^+/*Bdh1-Tfrc*^ mice. **(a)** Female B6.Del16^+/*Bdh1-Tfrc*^ mice displayed no significant differences in ambulatory activity compared to B6.WT littermates after saline administration (main effect of genotype, p>0.05). Female B6.WT mice displayed increased activity when given a 2.5mg/kg or a 7.5mg/kg dose of amphetamine compared to B6.WT littermates that received saline (main effect of treatment, p<0.0001). Female B6.Del16^+/*Bdh1-Tfrc*^ mice displayed attenuated activity following a 2.5mg/kg dose (treatment x genotype interaction, **p<0.005) and a 7.5mg/kg dose (treatment x genotype interaction, ****p<0.0001) of amphetamine compared to B6.WT littermates. **(b)** Male B6.Del16^+/*Bdh1-Tfrc*^ mice displayed no significant differences in ambulatory activity compared to B6.WT littermates after saline administration (main effect of genotype, p>0.05). Male B6.WT mice displayed increased activity when given a 2.5mg/kg or a 7.5mg/kg dose of amphetamine compared to B6.WT littermates that received saline (main effect of treatment, p<0.0001). Male B6.Del16^+/*Bdh1-Tfrc*^ mice displayed no significant changes in activity following a 2.5mg/kg dose (treatment x genotype interaction, p>0.05), but did show attenuated activity following a 7.5mg/kg dose (treatment x genotype interaction, ****p<0.0001) of amphetamine compared to B6.WT littermates. **(c)** Female B6.*Dlg1*^+/-^ mice displayed no significant differences in ambulatory activity compared to B6.WT littermates after saline administration (main effect of genotype, p>0.05). Female B6.WT mice displayed increased activity when given a 2.5mg/kg or a 7.5mg/kg dose of amphetamine compared to B6.WT littermates that received saline (main effect of treatment, p<0.0001). Female B6.*Dlg1*^+/-^ mice displayed no changes in activity after a 2.5mg/kg dose (treatment x genotype interaction, p>0.05) or a 7.5mg/kg dose (treatment x genotype interaction, p>0.05) compared to B6.WT littermates. **(d)** Male B6.*Dlg1*^+/-^ mice displayed no significant differences in ambulatory activity compared to B6.WT littermates after saline administration (main effect of genotype, p>0.05). Male B6.WT mice displayed increased activity when given a 2.5mg/kg or a 7.5mg/kg dose of amphetamine compared to B6.WT littermates that received saline (main effect of treatment, p<0.0001). Male B6.*Dlg1*^+/-^ mice displayed increased activity following a 2.5mg/kg dose (treatment x genotype interaction, ***p<0.001), but not a 7.5mg/kg dose (treatment x genotype interaction, p>0.05) of amphetamine compared to B6.WT littermates. All data were analyzed as outlined in methods. Results represent mean ± SEM (*p<0.05). B6.Del16^+/*Bdh1-Tfrc*^ mice-females: N=16 wild-type, 16 mutant; males: N=14 wild-type, 15 mutant. B6.*Dlg1*^+/-^ mice-females: N=8 wild-type, 10-11 mutant; males: 8-9 wild-type, 9 mutant.

### Anxiety-like behavior and associative memory are not changed in B6.Del16^+/*Bdh1-Tfrc*^ and B6.*Dlg1*^+/-^

We assessed performance in several tests for anxiety-like behavior (elevated plus maze, open field, marble burying) and did not observe any differences in the respective tasks (p>0.05) in B6.Del16^+/*Bdh1-Tfrc*^ or B6.*Dlg1*^+/-^ mice compared to B6.WT littermates **(Supplemental Figure 7) [statistics located in respective figure legend].**

We assessed associative learning and memory using context and cued fear conditioning. During training, female B6.Del16^+/*Bdh1-Tfrc*^ mice did not show any genotype-wide differences compared to B6.WT littermates, but did show a significant interaction (main effect of time x genotype, *F_21,630_*, p<0.0005). Sidak’s post-hoc analysis revealed female B6.Del16^+/*Bdh1-Tfrc*^ mice froze significantly more at the 300 (p<0.01), 360 (p<0.05), and 380 (p<0.0001) second time points compared to B6.WT littermates **(Supplemental Figure 8a)**. This result is consistent with the increased acoustic startle phenotype and suggests that female B6.Del16^+/*Bdh1-Tfrc*^ mice are more responsive to the immediate physiological and/or psychological effects of aversive stimuli. During the context (main effect of genotype, F_1,30_=0.1885, p>0.05) and cue (main effect of genotype, F_1,30_=2.682, p>0.05) tests, female B6.Del16^+/*Bdh1-Tfrc*^ mice had similar freezing percentages compared to B6.WT littermates **(Supplemental Figure 8b-c)**. Male B6.Del16^+/*Bdh1-Tfrc*^ mice did not show any differences during the training (main effect of genotype, *F_1,28_*=0.248, p>0.05), context (main effect of genotype, *F*_1,28_=0.9298, p>0.05), or cue (main effect of genotype, *F*_1,28_=1.981, p>0.05) phases compared to B6.WT littermates during any of the phases **(Supplemental Figure 8d-f)**.

During training, female B6.*Dlg1*^+/-^ mice froze significantly less (main effect of genotype, *F_1,17_*=5.096, p<0.05; main effect of genotype x time, *F_21,357_*=2.006, p<0.01) compared to B6.WT littermates **(Supplemental Figure 9a).** Sidak’s post-hoc analysis revealed B6.*Dlg1*^+/-^ females froze significantly less at 320s (p<0.01) and 380s (p<0.001). In the context test, female B6.*Dlg1*^+/-^ had similar freezing percentages compared to B6.WT littermates (main effect of genotype, *F_1,17_*=1.703, p>0.05) **(Supplemental Figure 9b)**, while female B6.*Dlg1*^+/-^ mice froze less (main effect of genotype, *F_1,17_*=4.302, p=0.05) during the cue test **(Supplemental Figure 9c)**. During training (main effect of genotype, *F_1,16_*=0.291, p>0.05), context (main effect of genotype, *F_1,16_*=0.4616, p>0.05) and cue tests (main effect of genotype, *F_1,16_*=0.007, p>0.05), male B6.*Dlg1*^+/-^ mice have a similar freezing percentage compared to B6.WT littermates **(Supplemental Figure 9e-f)**.

Collectively, we did not observe fear-dependent memory impairments in B6.Del16^+/*Bdh1-Tfrc*^ mice.

## Discussion

Here we report engineering a 3q29 deletion mouse model, and show it displays growth and behavioral deficits relevant to human 3q29 deletion syndrome phenotypes. Both female and male B6.Del16^+/*Bdh1-Tfrc*^ mice weigh significantly less than their B6.WT littermates, over the course of postnatal development and into adulthood, consistent with a prior study of humans with 3q29 deletion syndrome (2). In contrast, female but not male B6.*Dlg1*^+/-^ mice weigh significantly less than their B6.WT littermates suggesting that haploinsufficiency of *Dlg1* may be a partial contributor to the growth phenotype in female B6.Del16^+/*Bdh1-Tfrc*^ mice. We found multiple behavioral differences in B6.Del16^+/*Bdh1-Tfrc*^ mice that are endophenotypes of the neuropsychiatric disorders observed in 3q29 deletion study subjects (2, 9, 12). Male B6.Del16^+/*Bdh1-Tfrc*^ mice demonstrated social and cognitive impairment in the three-chambered social interaction and MWM tests, respectively, consistent with ASD and intellectual disability seen in humans with 3q29 deletion syndrome. Female B6.Del16^+/*Bdh1-Tfrc*^ mice showed an increased startle response consistent with reported hypersensitivity to acoustic stimuli in ASD patients (29). Female and male B6.Del16^+/*Bdh1-Tfrc*^ mice showed attenuated sensitivity to amphetamine-induced locomotor activity at a high dose of drug, suggesting perturbations in the dopaminergic system and demonstrating a shared behavior phenotype. Disruptions in dopamine signaling have been shown to confer risk for both ASD and SZ (30, 31). We also demonstrated that B6.*Dlg1*^+/-^ mice did not share any of the behavioral phenotypes that are observed in B6.Del16^+/*Bdh1-Tfrc*^ mice. These data indicate that B6.Del16^+/*Bdh1-Tfrc*^ mice display 3q29-related phenotypes that are not due solely to haploinsufficiency of *Dlg1*.

The finding that the B6.Del16^+/*Bdh1-Tfrc*^ mouse recapitulates aspects of 3q29 deletion at the genetic and phenotypic levels strongly suggests that the genetic drivers are *within* the 3q29 interval. The major difference between the mouse and human intervals is that the syntenic regions are inverted. This inversion occurred within the great ape lineage, as chimpanzees and humans have the same (derived) orientation of the region, but rhesus macaques have the inverted (ancestral) orientation that is present in mice. A copy-number neutral inversion of the region has been identified in humans, with no apparent phenotype (32). This implies that the critical event for the phenotype is the deletion itself, not a misregulation of a gene outside the deletion region. The fact that all 21 genes in the human 3q29 region are conserved in mouse and single gene mutations of each are being analyzed on the same genetic background through the International Mouse Phenotyping Consortium argues the region is ripe for genetic dissection. The B6.Del16^+/*Bdh1-Tfrc*^ mouse provides an entry point for such genetic dissection and ultimately, understanding of the circuitry and molecular mechanisms underlying these phenotypes. Towards this goal, the behavioral deficits provide clues to which brain regions may be most affected by the 3q29 deletion: the MWM is a hippocampal-dependent task (33, 34), social interaction is a striatum-dependent task, (35), and startle response is dependent on the caudal pontine reticular nucleus (36). Thus, the observed deficits in B6.Del16^+/*Bdh1-Tfrc*^ mice that we observed in the MWM are potential disruptions in these respective brain regions. Such relationships provide a starting point for evaluating the circuitry that may be disrupted in the B6.Del16^+/*Bdh1-Tfrc*^ mice, which may also be relevant for human 3q29 deletion patients.

While this current work focused on specific symptoms exhibited by 3q29 deletion patients, the mice have the potential to inform other 3q29 deletion phenotypes. The ongoing Emory 3q29 Project (http://genome.emory.edu/3q29/) has an active component where human study subjects participate in a comprehensive set of clinical evaluations (37). Phenotypic data from the B6.Del16^+/*Bdh1-Tfrc*^ mice can inform human phenotyping protocols and downstream analyses. For example, we observed sex-specific differences in our behavioral assertation of the B6.Del16^+/*Bdh1-Tfrc*^ mice but only limited patient phenotypes have been evaluated in a sex-specific way. We suspect that the sex-based differences we observed in the mice are due to intrinsic biological differences between females and males. Certainly, sex-specific differences are exhibited in other neuropsychiatric diseases, notably ASD where four times as many males as females are affected (38–40). Though purely speculative in relation to the 3q29 deletion, our behavior data demonstrates differences between B6.Del16^+/*Bdh1-Tfrc*^ females and male mice, which could be due to sex-specific molecular phenotypes. While the formal possibility exists that the variability in estrous stage contributed to the sex differences we observed, we think this is an unlikely explanation. The females were group-housed, minimizing any such variation. Furthermore, male mice display as much variability in behavioral assays as female mice without estrous control (41). These sex-specific differences are significant as they underscore the importance of studying both sexes in mice and humans.

Our *Dlg1*^+/-^ analyses argue that haploinsufficiency of *Dlg1* alone is not sufficient to explain the phenotypes associated with the B6.Del16^+/*Bdh1-Tfrc*^ mice, and points to either a different gene or a combination of genes within the interval that may or may not include *Dlg1* driving phenotypes. These data highlight the power of our approach. Recently, a neuron-specific deletion of *Dlg1* was reported to produce cognitive deficits in the males and motor learning deficits in the females (17). While this work identified differentially expressed genes in the hippocampus, the transcriptional profile is likely much different when only one copy of *DLG1* is deleted as in 3q29 deletion patients. The tissue-specific complete deletion of *Dlg1* also focuses on the role in neurons, but other cell types, such as glia, may contribute to 3q29 deletion associated phenotypes. One potential modifier of DLG1 is PAK2. In *Drosophila*, single *pak*-/+ or *dlg*-/+ mutants showed no phenotype but compound *pak*-/+ *dlg*-/+ flies showed decreased number of neuromuscular boutons indicating a genetic interaction (19). While this interaction remains unexamined in mouse, male *Pak2*^+/-^ mice are impaired on several social tasks including the three-chamber social interaction protocol used in the present study (18). Diminished *Pak2* likely contributes to the social impairment we observed in the B6.Del16^+/*Bdh1-Tfrc*^ male mice.

Several behavioral assays performed in the present study did not reveal phenotypes in the B6.Del16^+/*Bdh1-Tfrc*^ mice. Based on the known manifestations of 3q29 deletion in humans, we hypothesized that some of these phenotypes would be apparent, such as anxiety (2). It is possible that more complex human life experiences such as early life stress contribute to these phenotypes. Alternatively, our mouse studies were conducted on the C57BL/6N inbred strain, which may have obscured modifying genetic elements. Thus, examining the Del16^+/*Bdh1-Tfrc*^ on different genetic backgrounds could reveal additional impairments. It is also entirely possible that the distinctions in phenotype reflect fundamentally different biology between mice and humans. Nonetheless, the B6.Del16^+/*Bdh1-Tfrc*^ mouse will be an important tool to move forward and explore these issues and to systematically generate sub-deletions to genetically dissect the regions of the 3q29 interval driving the observed phenotypes. This 3q29 deletion mouse provides an excellent tool to continue the quest of unraveling the puzzle of neuropsychiatric disorders.

## Acknowledgements

GJB, TC, JGM, STW and DW designed the research. TR, RP and RP performed research with help from GG, SG, UK, RP, TM and JPS. ME, TR and RP analyzed data. TC, JGM, TR and DW wrote the manuscript.

All authors provided edits and approved final manuscript.

The work was supported by R01GM097331 (TC, DW, STW), R56MH116994 (TC, JM, DW, STW) and R01MH110701 along with funds from the Department of Human Genetics at Emory.

This study was supported in part by the Mouse Transgenic and Gene Targeting Core (TMF) and the Rodent Behavioral Core, which are subsidized by the Emory University School of Medicine and are part of the Emory Integrated Core Facilities.

Additional support was provided by the Georgia Clinical & Translational Science Alliance of the National Institutes of Health under Award Number UL1TR002378. The content is solely the responsibility of the authors and does not necessarily reflect the official views of the National Institutes of Health.

We are grateful to Dr. Jeffrey Miner, Washington University in St Louis, for providing *Dlg1* mutant mice.

This study was also supported by the Emory Winship Research Pathology Core Lab

## Genotyping *Dlg1*^+/-^ mice

PCR was performed using genomic ear or tail DNA, and the following primer pair: Dlg1_F5-TCAGAGACCACAAGAGGCCATTGGATACTC and Dlg1_R5-ATGCTGACTGGAAGGACTGCTAGTCTTCAG. These primers generate a 580bp mutant band and a 1071bp wild-type band. PCR conditions were as follows: 94°C for 5 min, 35 cycles at 94°C for 30 s, 56.7°C for 60 s, and 72°C for 60 s, then 1 cycle at 72°C for 10 min.

## Southern Blot and PCR to detect the mouse 3q29 deletion

To generate the 3q29 deletion in the mouse, two CRISPR gRNAs were designed at syntenic loci in the mouse genome: TTCAGTGGTATGTAACCCCTGG at Chr. 16:31,369,117 (**GRCm38.p3**) and CCTGAGCTGATTGGACAACTAG at Chr.16:32,634,414 (**GRCm38.p3**). The Emory Transgenic and Gene Targeting core injected 50ng/ml of each gRNAs and 100ng/ml Cas9 RNA into single-cell C57BL/6N zygotes. Embryos were cultured overnight and transferred to pseudopregnant females.PCR was used to screen for the deletion, and Southern blot analysis was used to assess for genomic rearrangements. PCR was performed using genomic ear or tail DNA, and the following primer pair: Proximal Forward-CCCTCCTTCCTCAATCACTG and Distal Reverse-TGCCACTCTTCAGCTCATTG (these primers generate a ~400bp product). PCR primers used to detect the breakpoint on the undeleted 3q29 interval (wild-type allele): Proximal Forward-CCCTCCTTCCTCAATCACTG and Proximal Reverse-CCCATCATTGGAGGAAAAA (these primers generate a ~400bp product). PCR conditions were as follows: 94°C for 5 min, 30 cycles at 94°C for 30 s, 56.7°C for 30 s, and 72°C for 60 s, then 1 cycle at 72°C for 5 min. A 542bp DNA probe for Southern blot analysis was generated using the following primer pair: forward primer-ATTCAGGTCTTTAATGAGAACACAA and reverse primer-TGAATAGTGGCTCTGTCTGAAG. Southern blots were adapted from a protocol as previously described (1). 10μg genomic DNA was digested overnight using the FastDigest XbaI from ThermoFisher Scientific (catalog number: FD0684).

## PCR Products for founders 127 and 131

Sequence of PCR products across the deletion that is shared between the two founders is underlined; sequence unique to each founder is in bold.

Founder 127:

> <u>AGGCATTTTCTCAAGTAAGGTTCCCTCTTTTCTGATGACTCTACCTTGTATCAAGTTGACATAAAGCTGCCAAGATCTGTCACCAGCTTCAGT</u>**GGTATGTAGGAGACAGACAGG**<u>AGACAGAGTTCCCTCCCTCTCCCTCCTGTCTGTCTTCTGTTCCTCTTTTATGTAGCAAACGTGACTCAGTGGCACGCCTCTCTTGCACTCCTATGAGATATCACTGAAATTATTATTATTATTATAAAAAAGAGAAACCGCCTCACTATTGGTGCCAAGAAAGGATTTTGGTGTCTAAGCATCTGGCCTCTGGGAACCAATGAG</u>

Founder 131:

> <u>AGGCATTTTCTCAAGTAAGGTTCCCTCTTTTCTGATGACTCTACCTTGTATCAAGTTGACATAAAGCTGCCAAGATCTGTCACCAGCTTCAGT</u>**GA**<u>AGACAGAGTTCCCTCCCTCTCCCTCCTGTCAAGCTGCCAAGATCTGTCACCAGCTTCAGTGAAGACAGAGTTCCCTCCCTCTCCCTCCTGTCTGTCTTCTGTTCCTCTTTTATGTAGCAAACGTGACTCAGTGGCACGCCTCTCTTGCACTCCTATGAGATATCACTGAAATTATTATTATTATTATAAAAAAGAGAAACCGCCTCACTATTGGTGCCAAGAAAGGATTTTGGTGTCTAAGCATCTGGCCTCTGGGAACCAATGAG</u>

The differences in sequence between the two founders is likely the of result of non-homologous end joining (NHEJ) commonly observed after CRISPR-mediated double-strand break repair (2)

## Western Blots

For Western blotting, brain tissue was disrupted in a Dounce homogenizer in freshly made, ice-cold buffer (20mM Tris-HCl pH 7.5, 118 mM NaCl, 4.7 mM KCl, 1.2 mM MgSO4, 2.5 mM CaCl2, 1.53 mM KH2PO4, 212.7 mM glucose, HALT Protease inhibitor (Thermo)). Samples were pelleted at 1000 x *g* for 20 minutes (at 4C) and sonicated in lysis buffer (50mM Tris-HCl pH 7.5, 40 mM NaCl, 1 mM EDTA, 0.5% Triton X-100, HALT Protease Inhibitor). Protein concentrations were normalized by BCA (ThermoFisher), reduced in Laemmli buffer and boiled at 95C. Protein samples (20ug) were loaded to each lane of 4-20% Tris-glycine pre-cast gels (Bio-rad) and transferred to nitrocellulose membranes using the TransBlot Turbo (Bio-rad). Membranes were blocked in Li-Cor blocking buffer and incubated with primary and secondary antibodies in 50:50 mixture of blocking buffer and PBS+0.1% Tween-20. Bands were visualized with fluorescent secondary antibodies (Li-Cor) in a Bio-rad ChemiDoc MP Imaging System, and band intensity was quantified using Image Lab software (Bio-rad). Each blot was normalized to a loaded control (Actin, Gapdh, or Tubulin depending on the size the protein of interest and source species of antibody). See **Supplemental Table 2** list of antibodies.

## Acoustic Startle and Growth Curve Analyses

### Acoustic Startle

Startle response to 70 Db was excluded from the dataset for all animals. Because the data were not normally distributed, the inverse normal function was used to transform the data to an approximately normal distribution. Proper transformation of the data was confirmed with the Shapiro-Wilk normality test implemented using the stats package (3). Linear mixed-effects models with restricted maximum likelihood estimation were implemented using the lme4 package (4). All models included decibel level as a fixed effect and subject ID as a random effect; males and females were analyzed separately and together. P values were calculated using Satterthwaite’s method using the lmerTest package (5).

### Growth Curves

All analyses were performed in R (3). We used the R package geepack to implement generalized estimating equations (GEE) that regressed weight measurements on genotype and age while accounting for within-subject correlation of measurements resulting from multiple time points of data collection (6–8). We analyzed males and females separately. We also repeated our analyses applying an inverse-normal transformation to the weight data to better satisfy modeling assumptions. Results using the raw and transformed data led to identical conclusions so we only present results from the analysis of the raw weight data in subsequent sections.

Using the GEE framework, we performed three distinct sets of analyses. We first compared weight measurements between B6.Del16^+/Bdh1-Tfrc^ mice and controls. We then compared weight measurements between B6.Dlg1^+/-^ mice and an independent set of controls. Finally, to see if the effect size of the B6.Del16^+/Bdh1-Tfrc^ genotype on weight differed in magnitude from the effect size for the B6.Dlg1^+/-^ genotype, we pooled the two datasets together and fit an additional GEE model that regressed weight measurements on genotype, age, dataset membership, and a genotype-by-dataset membership interaction term. We then tested the genotype-by-dataset interaction term to assess whether the effect size of the B6.Del16^+/Bdh1-Tfrc^ genotype significantly differed from the B6.Dlg1^+/-^ genotype.

## Additional Behavior Assays

### Circadian Rhythm

A mouse was placed into a plexiglass cage with the following dimensions: 48×25×22cm. Each cage was stocked with corncob bedding, food, and water for the duration of the assay. The plexiglass cage (locomotor chamber) was then placed between an apparatus that consisted of infrared breams (8×4 arrangement of beams, each beam is 5cm apart). When a mouse crossed broke two consecutive beams, it was considered one ambulation. Each mouse was in the locomotor chamber for 23 hours starting late morning. An attached computer collected the total number of ambulations, and the ambulations were binned per hour for each mouse.

### Elevated plus maze

The plus-shaped metal apparatus consisted of the following parameters: height of maze=81cm, open arms=30×8cm, closed arms=28×6cm, closed arm walls=17cm, and center region=8×6cm. Each mouse was placed in the center region of the maze, and was given 5 minutes to freely explore. The time spent in the open and closed portions of the maze was measured using CleverSystem’s (CleverSys Inc.) TopScan software.

### Open Field

This assay was run using an open field box with the following dimensions: 40×40×35cm. Mice were placed in the middle of the apparatus, and given 5 minutes to freely explore the arena. The time spent in the middle was measured using CleverSystem’s (CleverSys Inc.) TopScan software.

### Marble Burying

A plexiglass cage (48×25×22cm) was filled with ~2 inches of corncob bedding. The bedding was packed-down such that the surface was uniform. On top of the bedding, we placed 20 marbles (5 down, 4 across). A mouse was then placed into the prepared chamber and given 30 minutes to explore and bury the marble in the cage. After the 30 minutes, the mouse was carefully removed from the chamber. A picture of the chamber was then taken to assess how many marbles were buried. A marble was considered buried if it was >50% covered in bedding. The experimenter performing the scoring was blind to genotype.

### Social Interaction

We constructed a three-chamber apparatus with the following dimensions: 58×46×38cm. The north and south chambers had a length of 20cm while the middle chamber had a length of 18cm. Briefly, the subject mouse was acclimated to the middle portion of the three-chamber arena for 8 min followed by 10 minutes to acclimate to the entire arena. The subject mouse was enclosed in the middle region using doors that were manually inserted and removed at the appropriate time. Following acclimation, the subject mouse was then enclosed in the middle chamber, and an empty cup and a cup with a target mouse inside were placed in either the north or south chambers of the arena (alternated location of empty cup/cup with a mouse). The subject mouse was then given 10 min to interact with either the empty cup or cup with the target mouse. The final 10 min were video recorded, and time engaged in olfactory investigation of the empty cup and the cup with target mouse were recorded and scored by an experimenter blinded to genotype.

### Prepulse Inhibition

All mice were subjected to a series of increasing startle tones (75, 80, 85, 90, 95, 100, 105, 115, and 120db) for 40ms, and the response of the animal was measured by the instrument’s accelerometer. A startle curve was generated to ensure the subject mouse was responding to the stimulus. On day two, the animals were exposed to 6 blocks of startle conditions with each block consisting of 12 different trials; each trial is presented to the mice 6 times. The 12 different trials were presented randomly in each block and the animal’s response was measured after each trial. The 12 trials consisted of the following conditions: background (68db) for 20ms, Startle (120db) for 40ms, prepulses 1-4 (PP1=74db, PP2=78db, PP3=82db, and PP4=86db) for 20ms, and the prepulse-startle combinations (PP1.startle, PP2.startle, PP3.startle, and PP4.startle. During the prepulse-startle trials, the mice were exposed to the prepulse for 20ms and the startle for 40ms with a 100ms gap between the two.

### Fear Conditioning

To assess for deficits in associative learning and memory, we utilized a 3-day fear conditioning paradigm. On day 1, animals received 3 tone-shock pairings in a fear conditioning chamber where a tone was played for 20 sec followed by a 1 sec 0.5 milliamp shock (training). On day 2, animals were placed back into the same chamber for 7 min and time spent freezing was recorded (contextual memory). On day 3, mice were placed in a novel context and exposed to the shock-associated tone for 320 sec, and time spent freezing was recorded (cued memory).

## Histology

### Cresyl Violet and TUNEL Stain

Brains were sectioned on a cryostat at a thickness of 40μM. Sections were then placed through a series of ethanol washes for 5 minutes each: 100, 95, and 70%. After rinsing with distilled water, sections were then immersed into cresyl violet for 1 minute. Sections were again rinsed in distilled water before going through another series of 5-minute ethanol washes: 70 (5 drops of glacial acetic acid was only added to this ethanol), 95, and 100%. Sections were then cleared in xylene for 5 minutes followed by another 5-minute wash in fresh xylene

---

**Supplemental Table 1.**
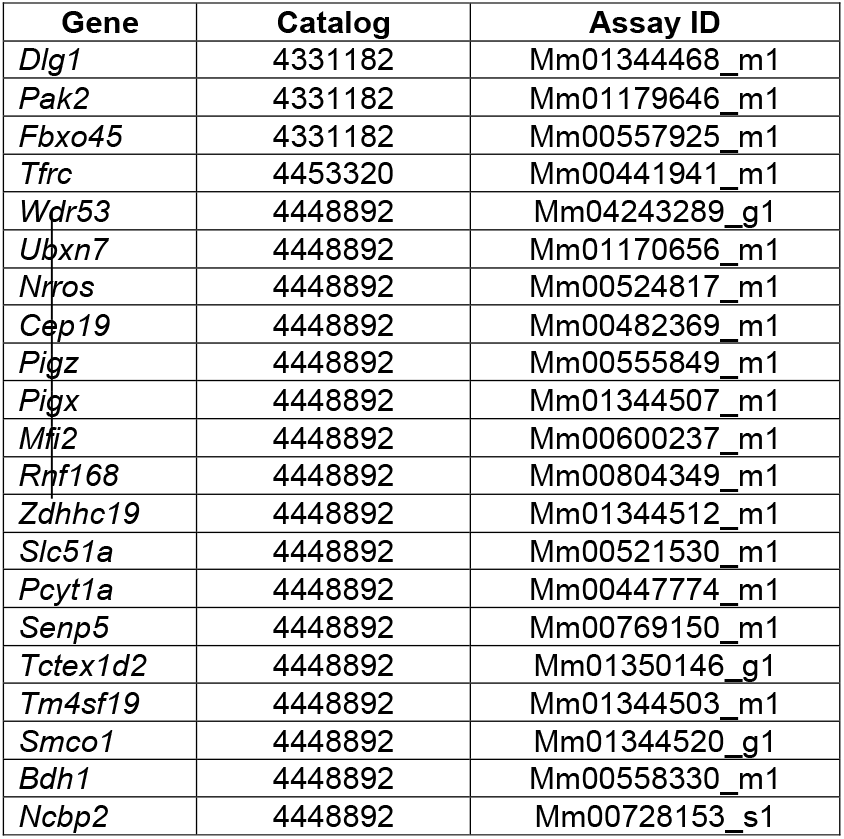

**Supplemental Table 2.**

| **Target** | **Supplier** | **Catalog Number** | **WB Dilution** |
| --- | --- | --- | --- |
| DLG1/SAP97 | Enzo | ADI-VAM-PS005-D | 1:1,000 |
| FBXO45 | Thermo Scientific | PA5-49458 | 1:2,000 |
| TFRC | Thermo Scientific | 13-6800 | 1:2,000 |
| PAK2 | Abcam | ab76293 | 1:2,000 |
| WDR53 | Novus Biological | NBP1-82758 | 1:250-500 |
| UBXN7 | Atlas Antibodies | HPA049442 | 1:1,000 |
| NCBP2 | Bethyl Laboratories | A302-553A | 1:1,000 |
| RNF168 | Novus Biological | AF7217 | 1:2,000 |
| PCYT1A | Abcam | Ab109263 | 1:2,000 |
| SENP5 | AAT Bioquest | 8C0368 | 1:1000 |
| BDH1 | Protein Tech | 15417-1-AP | 1:1000 |
| ACTIN | Thermo Scientific | AM4302 | 1:5,000 |

## Supplemental Figure Legends

**Supplemental Figure 1:**
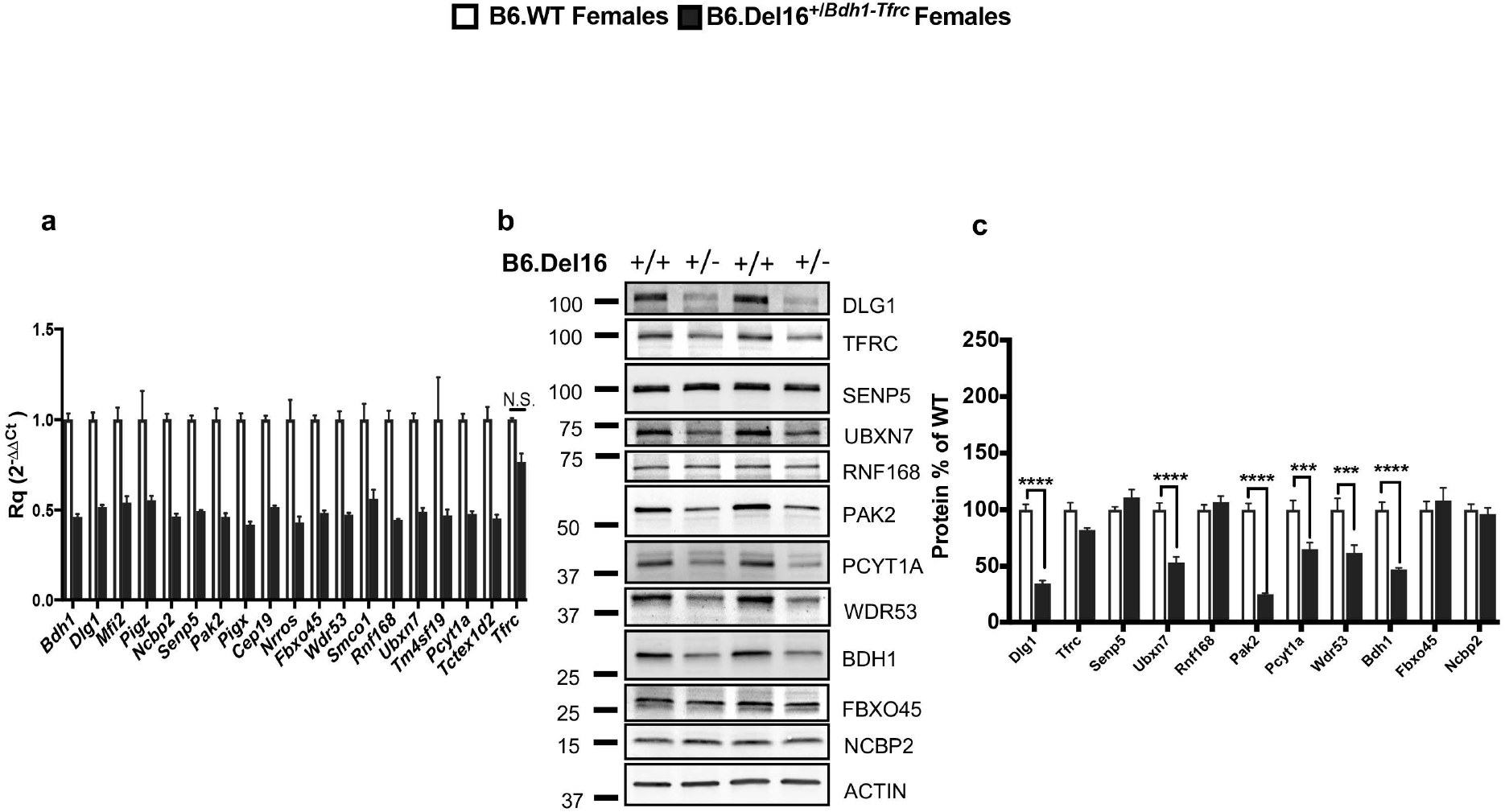
3q29-region specific gene and protein expression are reduced in the adult brains of B6.De16^+/*Bdh1-Tfrc*^ mice. **(a)** Gene expression analyses reveals most of the genes (with the exception of *Tfrc*) within the mouse 3q29 interval are ~50% reduced in both females. All genes, except *Tfrc*, reached significance (p<0.0001) (N=4 wild-type, 4 mutant; 3 technical replicates per sample). **(b)** Protein expression analyses reveals a significant reduction in DLG1, UBXN7, PAK2, PCYT1A, WDR53, and BDH1 (***p<0.001, ****p<0.0001) (N=5 wild-type, 6 mutant). All data analyzed by two-way, repeated measures Anova followed by multiple comparisons with Sidak’s correction. Results represent mean ± SEM.

**Supplemental Figure 2:**
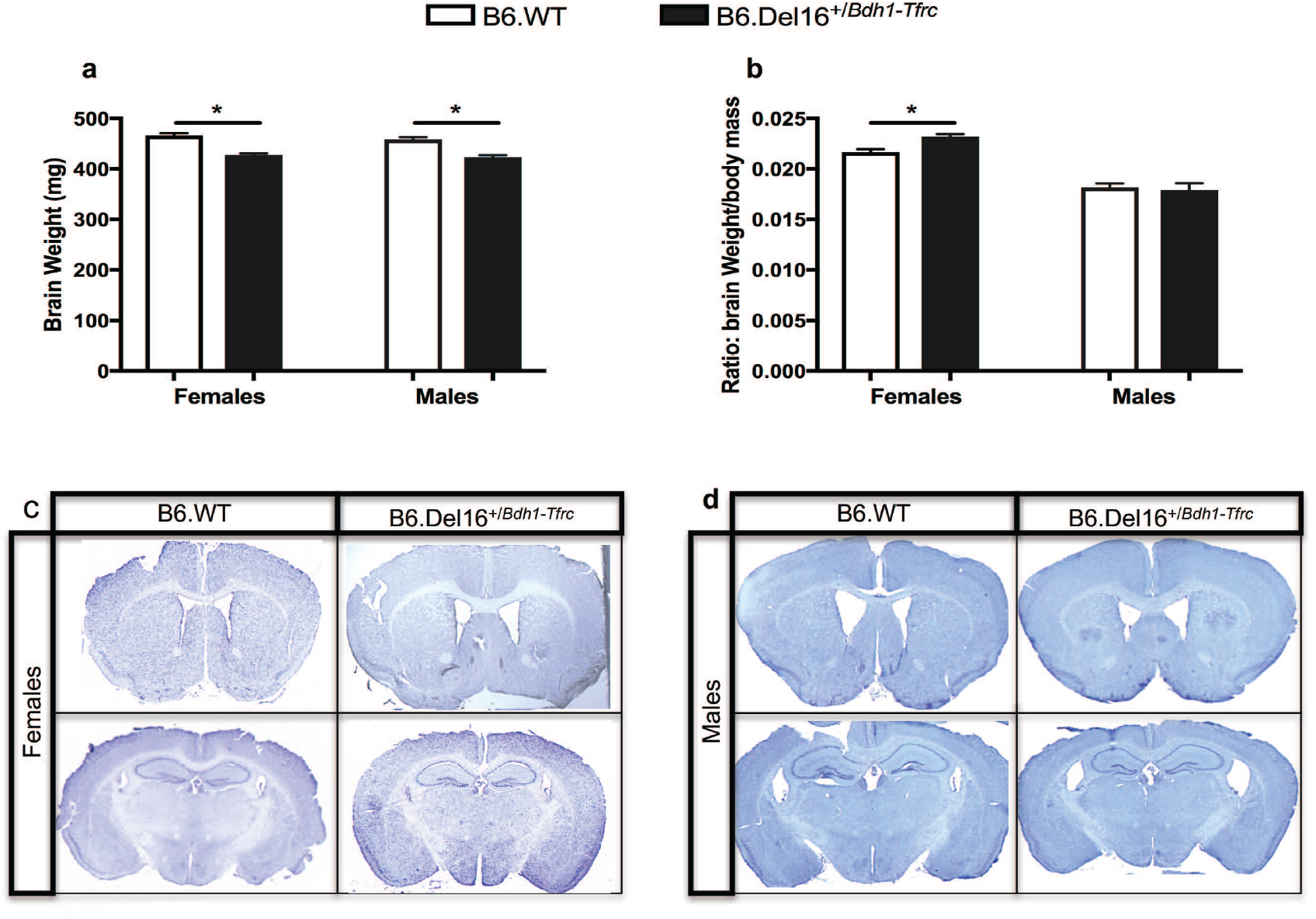
Size differences, but no gross phenotypes, in B6.Del16^+/*Bdh1-Tfrc*^ adult brains. **(a)** Female (****p<0.0001) [N=9 wild-type, 14 mutant] and male (***p<0.0005) [N=11 wild-type, 6 mutant] B6.De16^+/*Bdh1-Tfrc*^ mice have smaller brains compared to B6.WT littermates. **(b)** Female (**p<0.001) but not male (p>0.05) B6.De16^+/*Bdh1-Tfrc*^ mice have a larger brain weight:body mass ratio compared to B6.WT littermates. **(c)** Coronal sections of female B6.De16^+/*Bdh1-Tfrc*^ brains have normal morphology as shown via cresyl violet staining (N=4 wild-type, 4 mutant). **(d)** Coronal sections of male B6.De16^+/*Bdh1-Tfrc*^ brains have normal morphology as shown via cresyl violet staining (N=4 wild-type, 4 mutant). All data analyzed by unpaired, two-tailed t-test. Results represent mean ± SEM.

**Supplemental Figure 3:**
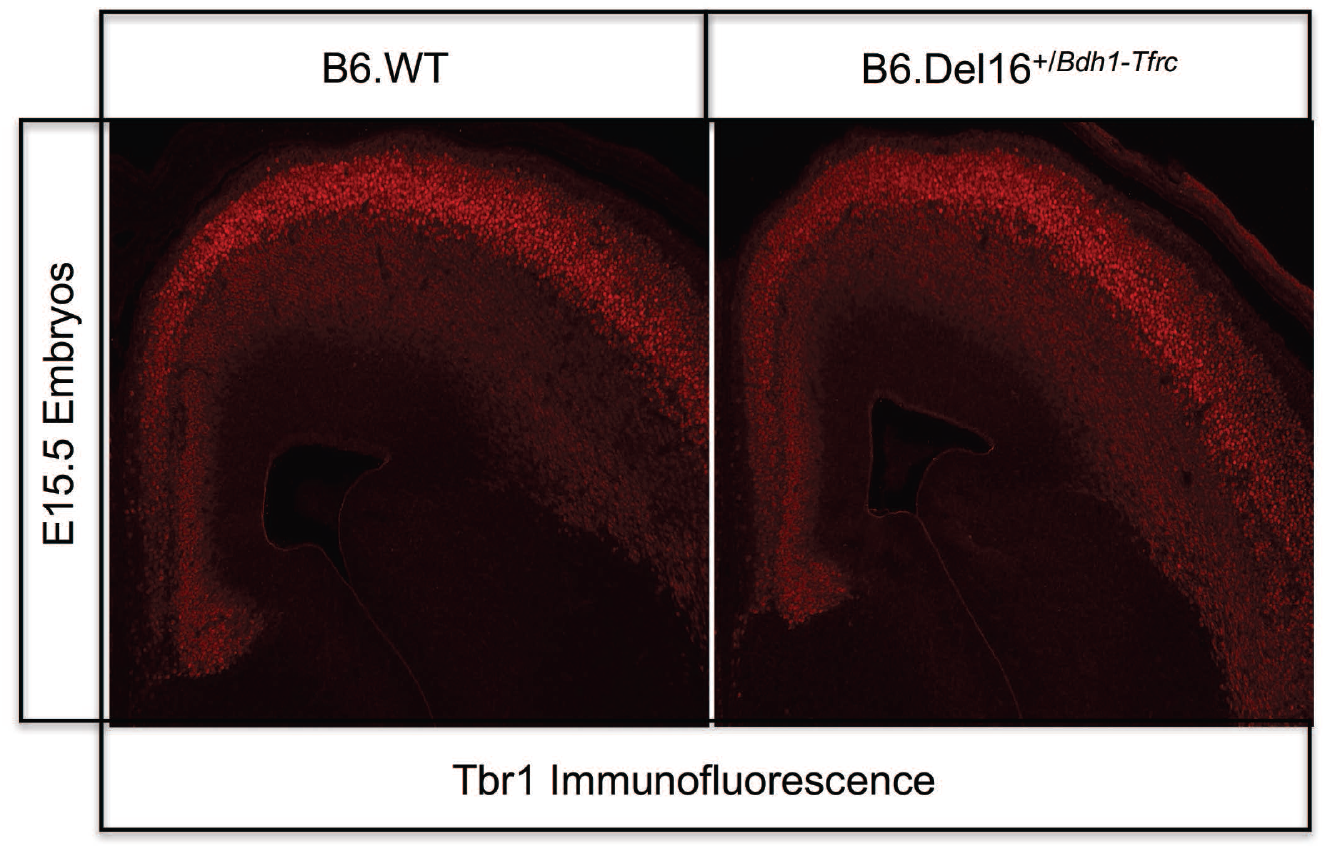
Cortical layer development appears normal in B6.De16^+/*Bdh1-Tfrc*^ embryos. E15.5 B6.De16^+/*Bdh1-Tfrc*^ embryos have similar TBR1 staining compared to B6.WT littermates (N=3 wild-type, 3 mutant).

**Supplemental Figure 4:**
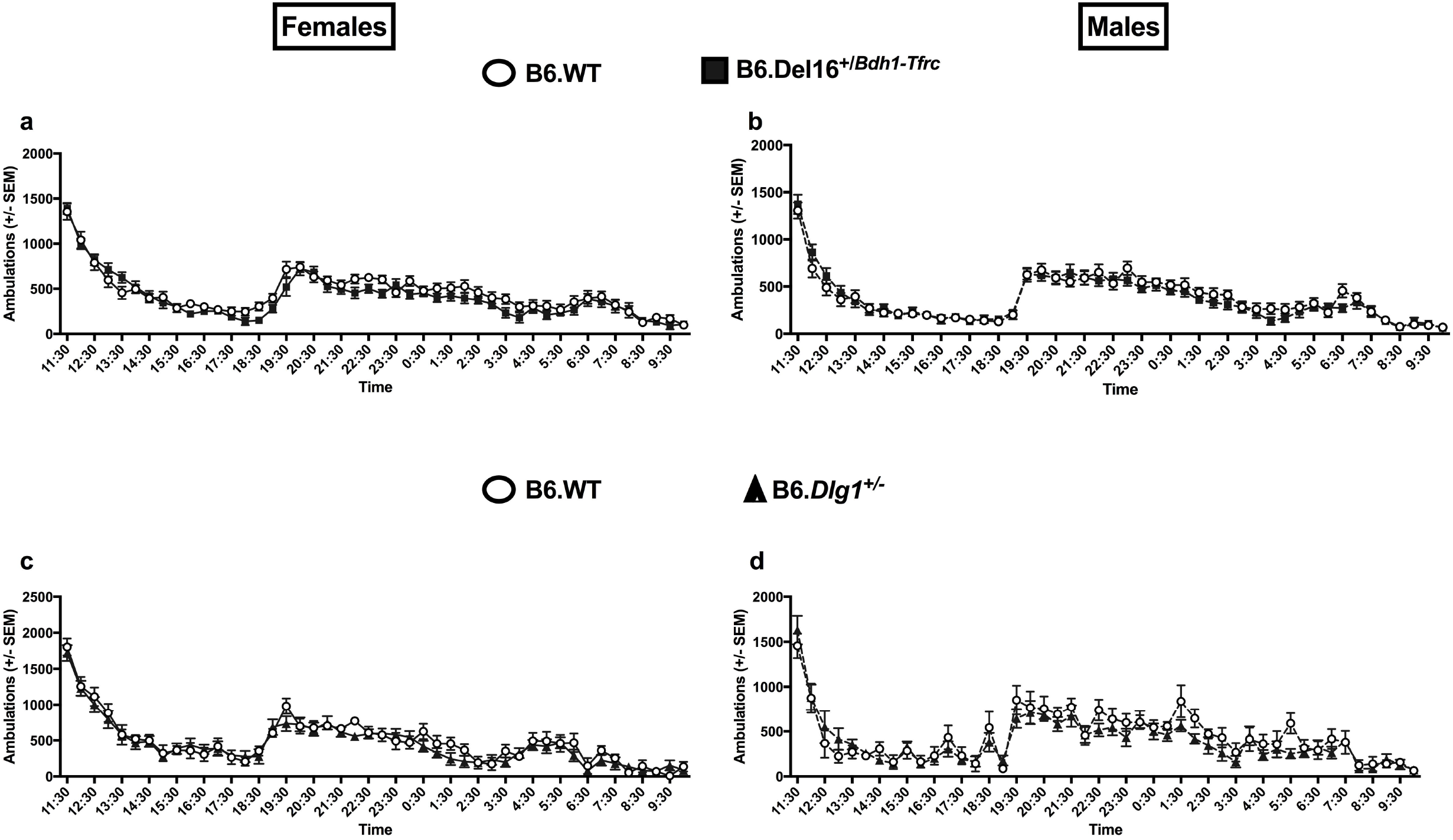
Locomotor activity is normal in both B6.Del16^+/*Bdh1-Tfrc*^ and B6.*Dlg1*^+/-^ mice. **(a)** Female (main effect of genotype, p>0.05) [N=16 wild-type. 16 mutant] and **(b)** male B6.De16^+/*Bdh1-Tfrc*^ (main effect of genotype, p>0.05) [N=15 wild-type, 15 mutant] display similar activity compared to their B6.WT littermates during the circadian rhythm paradigm. **(c)** Female (main effect of genotype, p>0.05) [N=7 wild-type, 10 mutant] and **(d)** male (main effect of genotype, p>0.05) [N=5 wild-type, 8 mutant] B6.*Dlg1*^+/-^ mice display similar activity compared to their B6.WT littermates during the circadian rhythm paradigm. All data analyzed by two-way, repeated measures Anova. Results represent mean ± SEM.

**Supplemental Figure 5:**
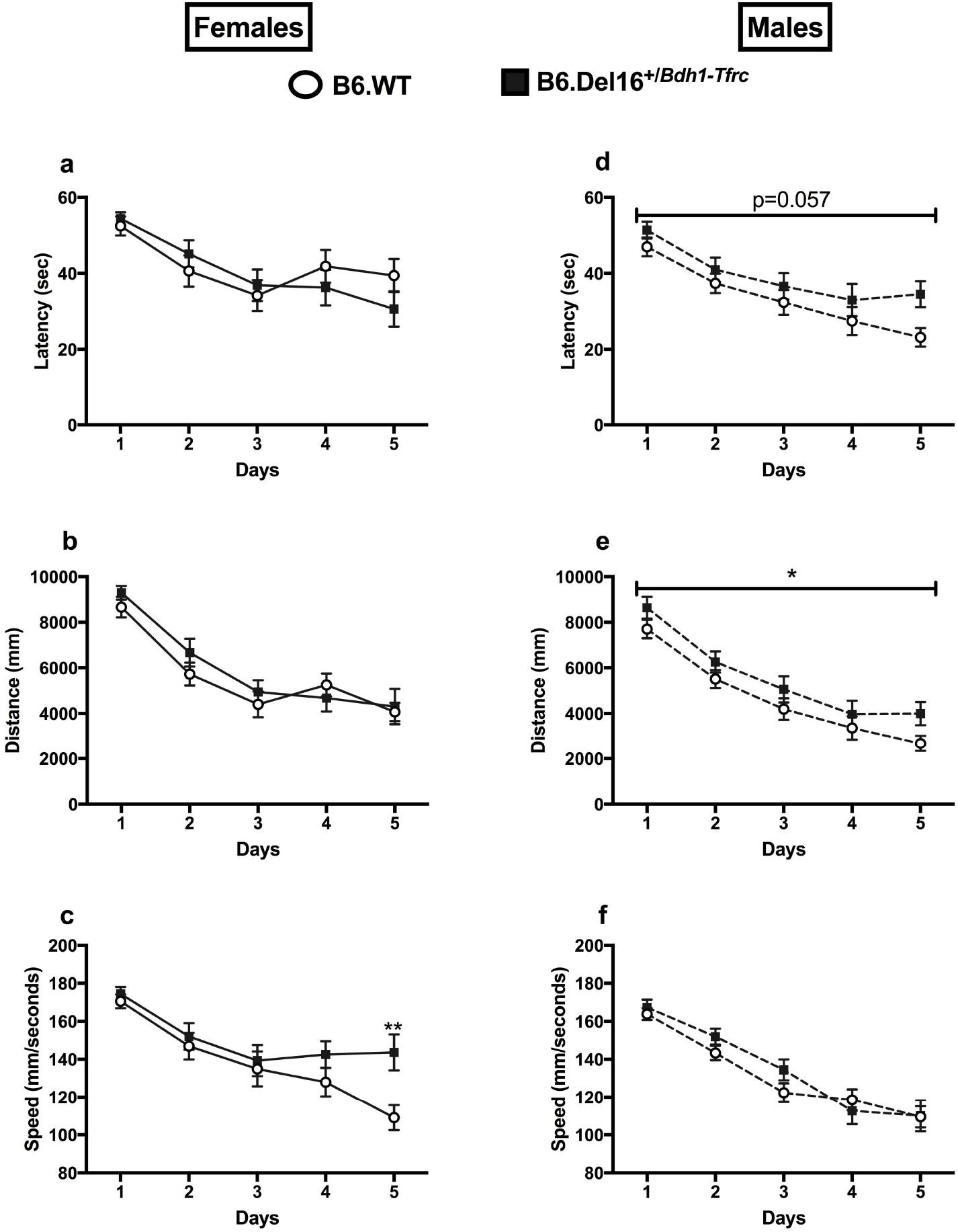
Sex-specific differences in B6.Del16^+/*Bdh1-Tfrc*^ mice during training portion of Morris water maze. **(a-c)** Female B6.De16^+/*Bdh1-Tfrc*^ mice show similar latency (main effect of genotype, p>0.05), distance (main effect of genotype, p>0.05), and swim speed (main effect of genotype, *F_1,30_*=2.188, p>0.05) compared to B6.WT littermates. There was a significant interaction (genotype x time, *F_4,120_*, p<0.005) for swim speed. Sidak’s post-hoc analysis revealed female B6.Del16^+/*Rdh1-Tfrc*^ mice swim faster on day 5 compared to B6.WT littermates (p<0.005) **(d-f)** Male B6.Del16^+/*Rdh1-Tfrc*^ mice show slightly elevated latency (main effect of genotype, p=0.057), distance (main effect of genotype, *p<0.05) but no differences in swim speed compared to their B6.WT littermates (main effect of genotype, p>0.05). All data analyzed by two-way, repeated measures Anova. Results represent mean ± SEM. Female mice: [N=16 wild-type, 16 mutant]; Male mice: [N=15 wild-type, 15 mutant].

**Supplemental Figure 6:**
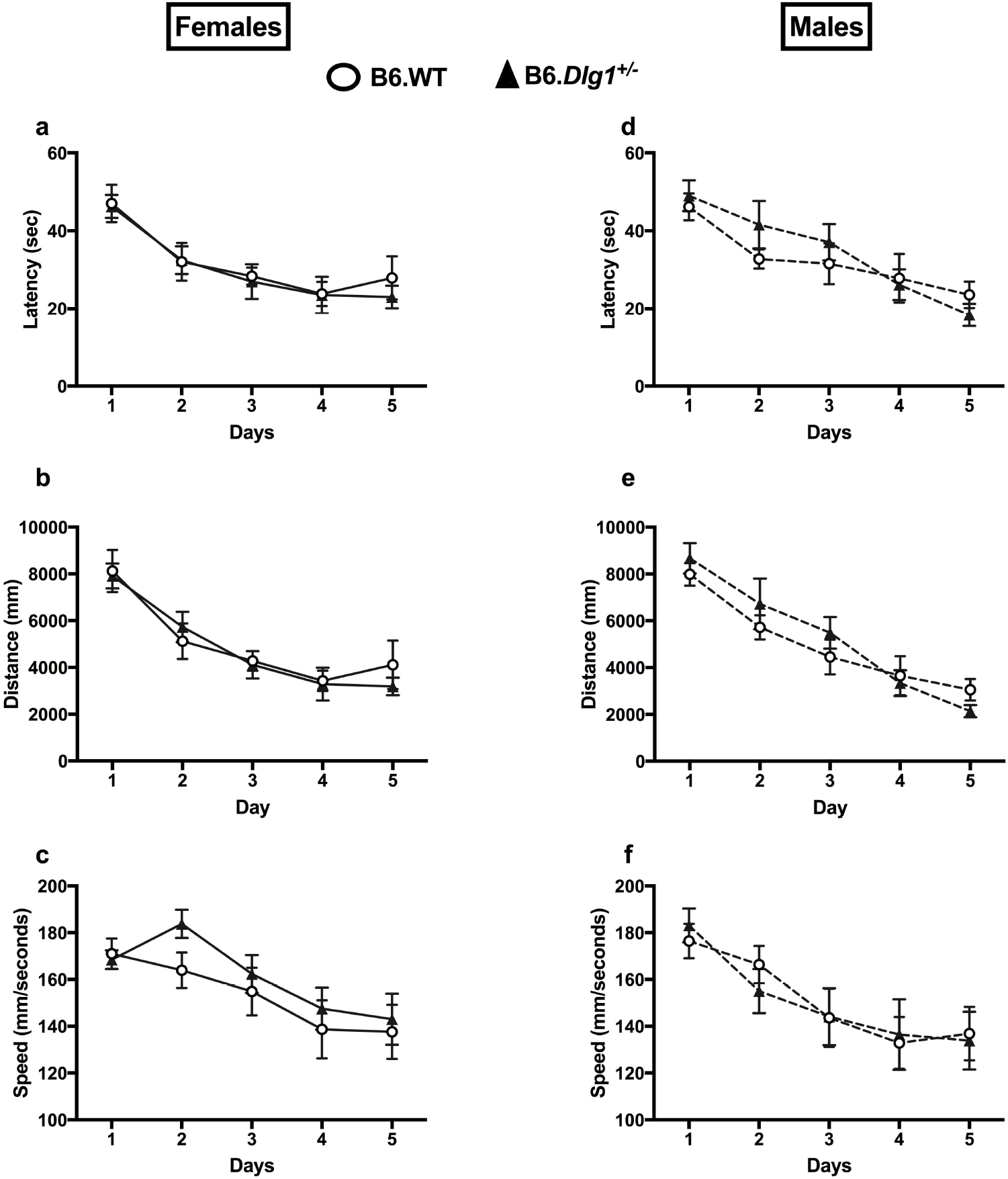
No differences in B6.Dlg1^+/-^ mice during training portion of Morris water maze. **(a-c)** Female B6.*Dlg1*^+/-^ mice show similar latencies (main effect of genotype, *F_1,17_*=0.1776, p>0.05), distance (main effect of genotype, *F_1,17_*=0.1407, p>0.05), and speed (main effect of genotype, *F_1,17_*=0.5809, p>0.05) compared to their B6.WT littermates. (d-f) Male B6.*Dlg1*^+/-^ mice show similar latency (main effect of genotype, *F_1,16_*=0.2816, p>0.05), distance (main effect of genotype, *F_1,16_*=0.3065, p>0.05), and swim speed (main effect of genotype, *F_1,16_*=0.0031, p>0.05) compared to their B6.WT littermates. All data analyzed by two-way, repeated measures Anova. Results represent mean ± SEM. Female mice: 8 wild-type, 11 mutant; Male mice: 9 wild-type, 9 mutant.

**Supplemental Figure 7:**
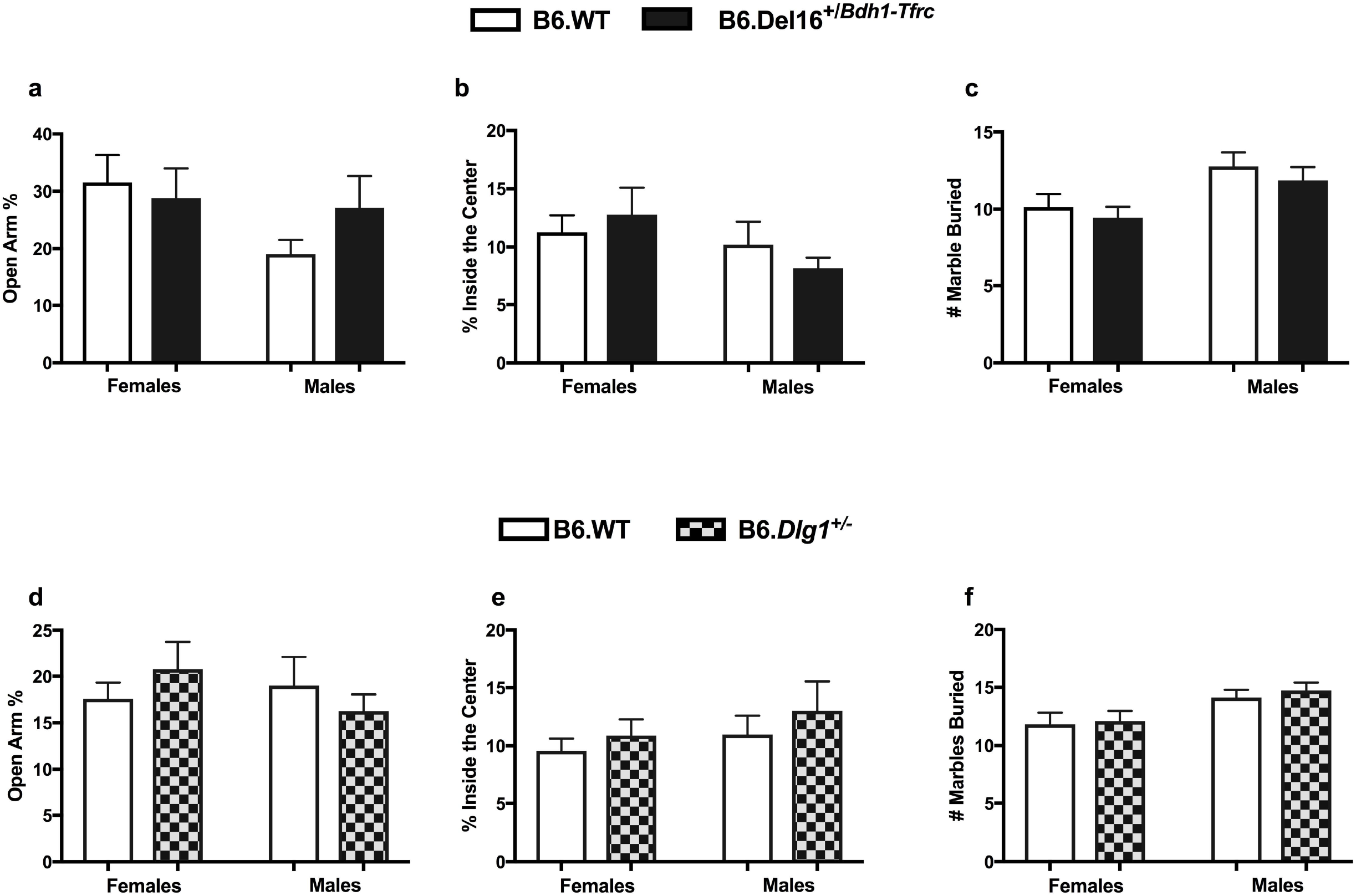
No anxiety-like behavior in neither B6.Del16^+/*Bdh1-Tfrc*^ nor B6.Dlg1^+/-^ mice. **(a)** Elevated Plus Maze: B6.Del16^+/*Rdh1-Tfrc*^ female ([N=16 wild-type, 16 mutant], t_30_=0.3795, p>0.05) and male ([N=15 wild-type, 15 mutant], t_28_=1 346, p>0.05) mice spend similar time on the open arm of the elevated plus maze compared to their B6.WT littermates. **(b)** Open Field: B6.Del16^+/*Bdh1-Tfrc*^ female ([N=16 wild-type, 16 mutant], t_30_=0.5561, p>0.05) and male ([N=15 wild-type, 15 mutant], t_28_=0.9245, p>0.05) mice spend similar time in the center of the open field compared to B6.WT littermates. (c) Marble Burying Task: B6.Del16^+/*Bdh1-Tfrc*^ female ([N=16 wild-type, 16 mutant], t_30_=0.6147, p>0.05) and male ([N=15 wild-type, 15 mutant], t_28_=0.7167, p>0.05) mice bury similar number of marbles compared to B6.WT littermates. **(d)** Elevated Plus Maze: B6.*Dlg1*^+/-^ female ([N=8 wild-type, 10 mutant], t_16_=0.8628, p>0.05) and male ([N=9 wild=type and 9 mutant], t_16_=0.7614, p>0.05) spend similar time on the open arm of the elevated plus maze compared to B6.WT littermates. **(e)** Open Field: B6.*Dlg1*^+/-^ female ([N=14 wild-type, 13 mutant], t_25_=0.7548, p>0.05) and male ([N=12 wild-type, 13 mutant], t_23_=0.667, p>0.05) mice spend similar time in the center of the open field compared to B6.WT littermates. **(f)** Marble Burying: B6.*Dlg1*^/-^ female ([N=14 wild-type, 13 mutant], t_25_=0.22, p>0.05) and male ([N=12 wild-type, 13 mutant], t_23_=0.632, p>0.05) mice bury similar number of marbles compared to B6.WT littermates. All data analyzed by two-tailed Students t-test. Results represent the mean ± SEM.

**Supplemental Figure 8:**
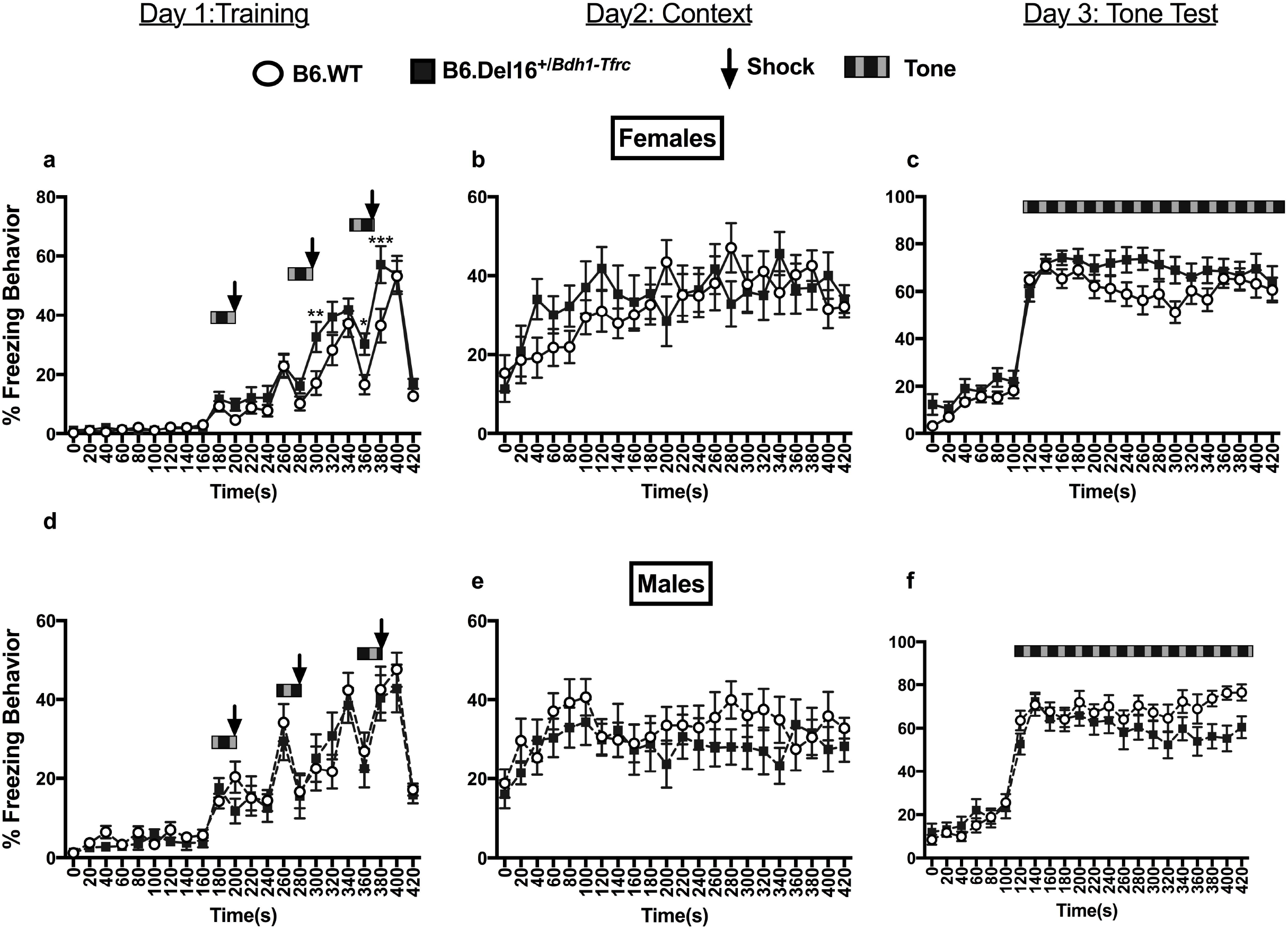
Associative learning and memory is not impaired in B6.Del16^+/*Bdh1-Tfrc*^ mice. **(a)** Female B6.Del16^+/*Bdh1-Tfrc*^ mice freeze at similar percentages during the training phase compared to B6.WT littermates, but there was a significant interaction (main effect of time, *F_21,630_*, p<0.0005). Sidak’s post-hoc analysis revealed significantly more freezing at the 300(** p<0.01), 360 (*p<0.05), and 380s (***p<0.0001) time points compared to B6.WT littermates **(b)** Female B6.Del16^+/*Bdh1-Tfrc*^ mice freeze at similar percentages during the contextual phase compared to B6.WT littermates (main effect of genotype, *F*_1,30_=0.1885, p>0.05). **(c)** Female B6.Del16^+/*Bdh1-Tfrc*^ mice freeze at similar percentages during the tone phase compared to B6.WT littermates (main effect of genotype, *F_1,30_*=2.682, p>0.05). (d-f) Male B6.*Del16*^+/*Bdh1-Tfrc*^ mice freeze at similar percentages during the training (main effect of genotype, *F_1,28_*=0.248, p>0.05), context (main effect of genotype, *F*_1,28_=0.9298, p>0.05), and cue (main effect of genotype, *F*_1,28_=1.981, p>0.05) phases compared to B6.WT littermates. All data analyzed by two-way, repeated measures Anova. Results represent mean ± SEM, Female mice: N=16 wild-type, 16 mutant. Male mice: N=15 wild-type, 15 mutant.

**Supplemental Figure 9:**
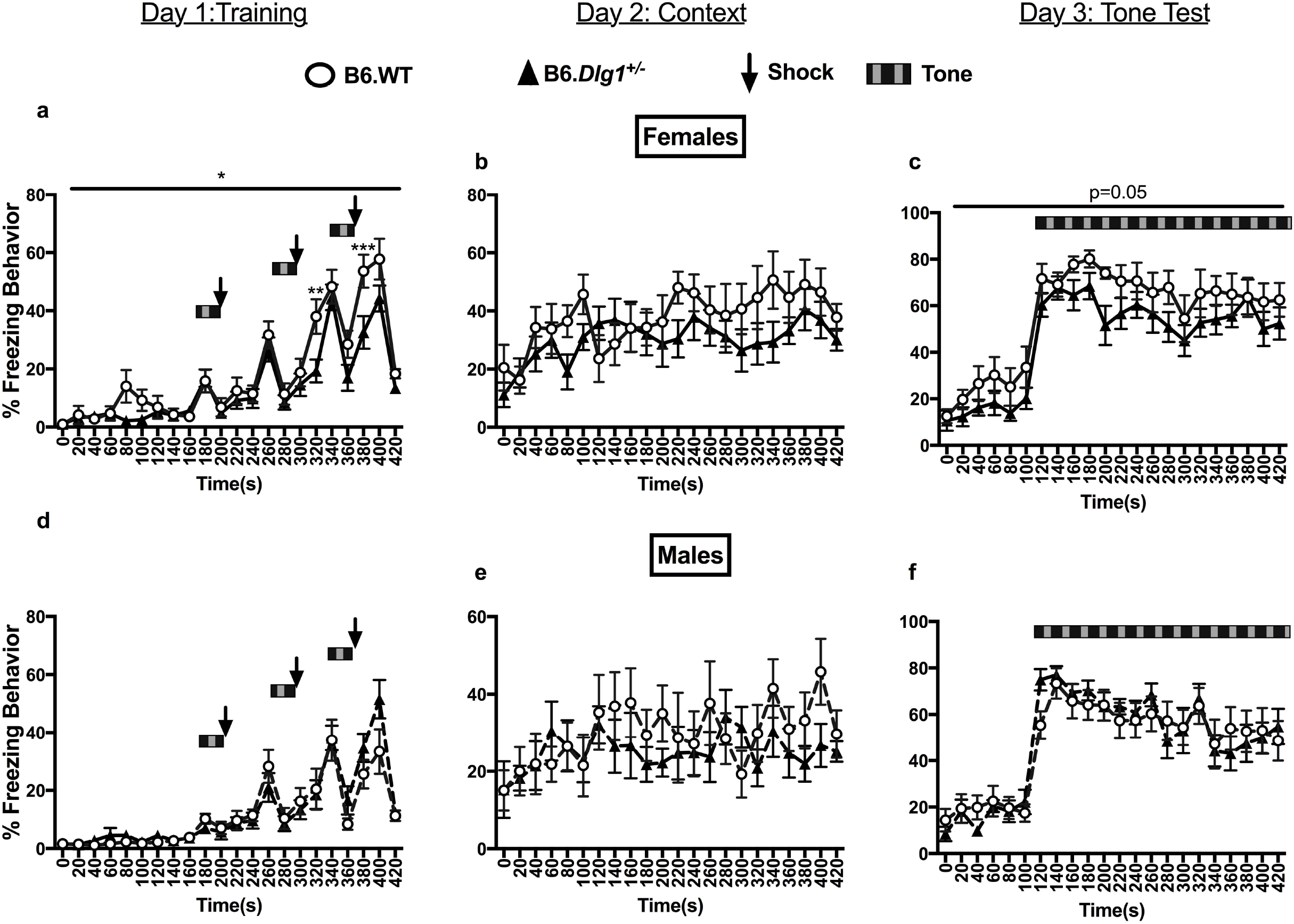
Associative learning and memory is not impaired in B6.*Dlg1*^+/-^ mice. (a) Female B6.*Dlg1*^+/-^ mice freeze less during the training phase compared to B6.WT littermates (main effect of genotype, *F_1,17_*=5.096, *p<0.05; main effect of time, F_21,357_=2.006, **p<0.01). Sidak’s post-hoc analysis revealed B6.*Dlg1*^+/-^ females froze significantly less at 320s (*p<0.05) and 380s (***p<0.001). **(b)** Female B6.*Dlg1*^+/-^ mice freeze at similar percentages during the contextual phase compared to B6.WT littermates (main effect of genotype, F_1,17_=1.703, p>0.05). (c) Female B6.*Dlg1*^+/-^ mice freeze slightly less during the tone phase compared to B6.WT littermates (main effect of genotype, *F_1,17_*=4.302, p=0.05) **(d-f)** Male B6.*Dlg1*^+/-^ mice freeze at similar percentages during the training (main effect of genotype, *F_1,16_*=0.291, p>0.05), context (main effect of genotype, *F_1,16_*=0.4616, p>0.05), and cue (main effect of genotype, *F*_1,16_=0.007, p>0.05) phases compared to B6.WT littermates. All data analyzed by two-way, repeated measures Anova. Results represent mean ± SEM. Female mice: N=8 wild-type, 11 mutant. Male mice: N=9 wild-type, 9 mutant.

## References

1. Ballif BC, Theisen A, Coppinger J, Gowans GC, Hersh JH, Madan-Khetarpal S, et al. Expanding the clinical phenotype of the 3q29 microdeletion syndrome and characterization of the reciprocal microduplication. Mol Cytogenet. 2008;1:8.

2. Glassford MR, Rosenfeld JA, Freedman AA, Zwick ME, Mulle JG, Unique Rare Chromosome Disorder Support G. Novel features of 3q29 deletion syndrome: Results from the 3q29 registry. Am J Med Genet A. 2016;170A(4):999–1006.

3. Marshall CR, Howrigan DP, Merico D, Thiruvahindrapuram B, Wu W, Greer DS, et al. Contribution of copy number variants to schizophrenia from a genome-wide study of 41,321 subjects. Nat Genet. 2017;49(1):27–35.

4. Mulle JG, Dodd AF, McGrath JA, Wolyniec PS, Mitchell AA, Shetty AC, et al. Microdeletions of 3q29 confer high risk for schizophrenia. Am J Hum Genet. 2010;87(2):229–36.

5. Mulle JG, Gambello MJ, Cook EH, Rutkowski TP, Glassford M. 3q29 Recurrent Deletion. In: Adam MP, Ardinger HH, Pagon RA, Wallace SE, Bean LJH, Stephens K, et al., editors. GeneReviews((R)). Seattle (WA)1993.

6. Sanders SJ, He X, Willsey AJ, Ercan-Sencicek AG, Samocha KE, Cicek AE, et al. Insights into Autism Spectrum Disorder Genomic Architecture and Biology from 71 Risk Loci. Neuron. 2015;87(6):1215–33.

7. Willatt L, Cox J, Barber J, Cabanas ED, Collins A, Donnai D, et al. 3q29 microdeletion syndrome: clinical and molecular characterization of a new syndrome. Am J Hum Genet. 2005;77(1):154–60.

8. Grozeva D, Conrad DF, Barnes CP, Hurles M, Owen MJ, O’Donovan MC, et al. Independent estimation of the frequency of rare CNVs in the UK population confirms their role in schizophrenia. Schizophr Res. 2012;135(1–3):1–7.

9. Mulle JG. The 3q29 deletion confers >40-fold increase in risk for schizophrenia. Mol Psychiatry. 2015;20(9):1028–9.

10. Korablev AN, Serova IA, Serov OL. Generation of megabase-scale deletions, inversions and duplications involving the Contactin-6 gene in mice by CRISPR/Cas9 technology. BMC Genet. 2017;18(Suppl 1):112.

11. Carroll LS, Williams HJ, Walters J, Kirov G, O’Donovan MC, Owen MJ. Mutation screening of the 3q29 microdeletion syndrome candidate genes DLG1 and PAK2 in schizophrenia. Am J Med Genet B Neuropsychiatr Genet. 2011;156B(7):844–9.

12. Rutkowski TP, Schroeder JP, Gafford GM, Warren ST, Weinshenker D, Caspary T, et al. Unraveling the genetic architecture of copy number variants associated with schizophrenia and other neuropsychiatric disorders. J Neurosci Res. 2017;95(5):1144–60.

13. Leonard AS, Davare MA, Horne MC, Garner CC, Hell JW. SAP97 is associated with the alpha-amino-3-hydroxy-5-methylisoxazole-4-propionic acid receptor GluR1 subunit. J Biol Chem. 1998;273(31):19518–24.

14. Purcell SM, Moran JL, Fromer M, Ruderfer D, Solovieff N, Roussos P, et al. A polygenic burden of rare disruptive mutations in schizophrenia. Nature. 2014;506(7487):185–90.

15. Uezato A, Kimura-Sato J, Yamamoto N, Iijima Y, Kunugi H, Nishikawa T. Further evidence for a male-selective genetic association of synapse-associated protein 97 (SAP97) gene with schizophrenia. Behav Brain Funct. 2012;8:2.

16. Toyooka K, Iritani S, Makifuchi T, Shirakawa O, Kitamura N, Maeda K, et al. Selective reduction of a PDZ protein, SAP-97, in the prefrontal cortex of patients with chronic schizophrenia. J Neurochem. 2002;83(4):797–806.

17. Gupta P, Uner OE, Nayak S, Grant GR, Kalb RG. SAP97 regulates behavior and expression of schizophrenia risk enriched gene sets in mouse hippocampus. PLoS One. 2018;13(7):e0200477.

18. Wang Y, Zeng C, Li J, Zhou Z, Ju X, Xia S, et al. PAK2 Haploinsufficiency Results in Synaptic Cytoskeleton Impairment and Autism-Related Behavior. Cell Rep. 2018;24(8):2029–41.

19. Grice SJ, Liu JL, Webber C. Synergistic interactions between Drosophila orthologues of genes spanned by de novo human CNVs support multiple-hit models of autism. PLoS Genet. 2015;11(3):e1004998.

20. Mahoney ZX, Sammut B, Xavier RJ, Cunningham J, Go G, Brim KL, et al. Discs-large homolog 1 regulates smooth muscle orientation in the mouse ureter. Proc Natl Acad Sci U S A. 2006;103(52):19872–7.

21. Yang M, Silverman JL, Crawley JN. Automated three-chambered social approach task for mice. Curr Protoc Neurosci. 2011;Chapter 8:Unit 8 26.

22. Chalermpalanupap T, Schroeder JP, Rorabaugh JM, Liles LC, Lah JJ, Levey AI, et al. Locus Coeruleus Ablation Exacerbates Cognitive Deficits, Neuropathology, and Lethality in P301S Tau Transgenic Mice. J Neurosci. 2018;38(1):74–92.

23. Weinshenker D, Miller NS, Blizinsky K, Laughlin ML, Palmiter RD. Mice with chronic norepinephrine deficiency resemble amphetamine-sensitized animals. Proc Natl Acad Sci U S A. 2002;99(21):13873–7.

24. R-CoreTeam. A language and environment for statistical computing. R foundation for Statistical Computing. Vienna, Austria2017. p. http://www.R-project.org.

25. Bates D, Machler M, Bolker BM, Walker SC. Fitting Linear Mixed-Effects Models Using lme4. J Stat Softw. 2015;67(1):1–48.

26. Kuznetsova A, Brockhoff PB, Christensen RHB. lmerTest Package: Tests in Linear Mixed Effects Models. J Stat Softw. 2017;82(13):1–26.

27. Cox DM, Butler MG. A clinical case report and literature review of the 3q29 microdeletion syndrome. Clin Dysmorphol. 2015;24(3):89–94.

28. Nielsen J, Fejgin K, Sotty F, Nielsen V, Mork A, Christoffersen CT, et al. A mouse model of the schizophrenia-associated 1q21.1 microdeletion syndrome exhibits altered mesolimbic dopamine transmission. Transl Psychiatry. 2017;7(11):1261.

29. Takahashi H, Nakamura T, Kim J, Kikuchi H, Nakahachi T, Ishitobi M, et al. Acoustic Hyper-Reactivity and Negatively Skewed Locomotor Activity in Children With Autism Spectrum Disorders: An Exploratory Study. Front Psychiatry. 2018;9:355.

30. Kesby JP, Eyles DW, McGrath JJ, Scott JG. Dopamine, psychosis and schizophrenia: the widening gap between basic and clinical neuroscience. Transl Psychiatry. 2018;8(1):30.

31. Paval D. A Dopamine Hypothesis of Autism Spectrum Disorder. Dev Neurosci. 2017;39(5):355–60.

32. Kidd JM, Cooper GM, Donahue WF, Hayden HS, Sampas N, Graves T, et al. Mapping and sequencing of structural variation from eight human genomes. Nature. 2008;453(7191):56–64.

33. Logue SF, Paylor R, Wehner JM. Hippocampal lesions cause learning deficits in inbred mice in the Morris water maze and conditioned-fear task. Behav Neurosci. 1997;111(1):104–13.

34. Morris RG, Hagan JJ, Rawlins JN. Allocentric spatial learning by hippocampectomised rats: a further test of the "spatial mapping" and "working memory" theories of hippocampal function. Q J Exp Psychol B. 1986;38(4):365–95.

35. Baez-Mendoza R, Schultz W. The role of the striatum in social behavior. Front Neurosci. 2013;7:233.

36. Koch M. The neurobiology of startle. Prog Neurobiol. 1999;59(2):107–28.

37. Murphy MM, Lindsey Burrell T, Cubells JF, Espana RA, Gambello MJ, Goines KCB, et al. Study protocol for The Emory 3q29 Project: evaluation of neurodevelopmental, psychiatric, and medical symptoms in 3q29 deletion syndrome. BMC Psychiatry. 2018;18(1):183.

38. Halladay AK, Bishop S, Constantino JN, Daniels AM, Koenig K, Palmer K, et al. Sex and gender differences in autism spectrum disorder: summarizing evidence gaps and identifying emerging areas of priority. Mol Autism. 2015;6:36.

39. Kirkovski M, Enticott PG, Fitzgerald PB. A review of the role of female gender in autism spectrum disorders. J Autism Dev Disord. 2013;43(11):2584–603.

40. Lai MC, Lombardo MV, Baron-Cohen S. Autism. Lancet. 2014;383(9920):896–910.

41. Prendergast BJ, Onishi KG, Zucker I. Female mice liberated for inclusion in neuroscience and biomedical research. Neurosci Biobehav Rev. 2014;40:1–5.

## References

1. Xie N, Gong H, Suhl JA, Chopra P, Wang T, Warren ST. Reactivation of FMR1 by CRISPR/Cas9-Mediated Deletion of the Expanded CGG-Repeat of the Fragile X Chromosome. PLoS One. 2016;11(10):e0165499.

2. Zhang JH, Adikaram P, Pandey M, Genis A, Simonds WF. Optimization of genome editing through CRISPR-Cas9 engineering. Bioengineered. 2016;7(3):166–74.

3. R-CoreTeam. A language and environment for statistical computing. R foundation for Statistical Computing. Vienna, Austria2017. p. http://www.R-project.org.

4. Bates D, Machler M, Bolker BM, Walker SC. Fitting Linear Mixed-Effects Models Using lme4. J Stat Softw. 2015;67(1):1–48.

5. Kuznetsova A, Brockhoff PB, Christensen RHB. lmerTest Package: Tests in Linear Mixed Effects Models. J Stat Softw. 2017;82(13):1–26.

6. Højsgaard S, Halekoh U, Yan J. The R Package geepack for Generalized Estimating Equations. J Stat Softw. 2006;15(2):1–11.

7. Yan J. geepack: Yet another package for generalized estimating. Equations R-News. 2002;2/3:12–4.

8. Yan J, Fine J. Estimating equations for association structures. Stat Med. 2004;23(6):859–74.

